# Nucleobase adduct-containing metabolites are MR1 ligands that stimulate self-reactive MR1T cells

**DOI:** 10.1101/2022.08.26.505411

**Authors:** Alessandro Vacchini, Qinmei Yang, Andrew Chancellor, Julian Spagnuolo, Daniel Joss, Ove Øyås, Aisha Beshirova, Corinne De Gregorio, Michael Pfeffer, Jörg Stelling, Daniel Häussinger, Marco Lepore, Lucia Mori, Gennaro De Libero

**Affiliations:** Experimental Immunology, Department of Biomedicine, University Hospital and University of Basel, 4031 Basel, Switzerland; Department of Chemistry, University of Basel, 4056 Basel, Switzerland; Department of Biosystems Science and Engineering and SIB Swiss Institute of Bioinformatics, ETH Zurich, 4058 Basel, Switzerland; Department of Chemistry, Biotechnology and Food Science, Norwegian University of Life Sciences, 1432 Ås, Norway

## Abstract

MR1T lymphocytes are a recently identified population of T cells that recognize unknown self-antigens presented by the non-polymorphic MHC-I-related molecule, MR1. MR1T cells can kill tumor cells and modulate the functions of other immune cells with promising therapeutic applications. By integrating genetic, pharmacological and biochemical approaches we identified carbonyl stress and alterations of nucleobase metabolism in tumor target cells that promote recognition by MR1 T cells. We dissected these pathways and found that nucleobase adduct-containing metabolites are self-antigens stimulating MR1T cells. Several nucleobase adducts are presented by MR1 molecules and stimulate individual MR1T cells. Our data suggest that MR1T cells are surveyor of cellular metabolic alterations occurring in conditions of metabolic stress, such as cancer, and lay the groundwork for the development of novel HLA-unrestricted T cell-based therapies.

## Introduction

MR1 is a non-polymorphic, ubiquitously expressed molecule that was initially characterized as the restriction element for mucosal-associated invariant-T (MAIT) cells (*1*), enabling MAIT cell detection of microbes during infection (*2-4*). The MAIT cell-stimulating microbial antigens were later identified as metabolites produced in the riboflavin pathway (*4, 5*). Subsequent studies revealed that the MR1 antigen-binding pocket can accommodate various metabolites and chemical entities (*4-8*). Accordingly, parallel studies revealed an additional population of MR1-restricted T cells, named MR1T cells, that do not respond to microbial antigens while reacting at clonal level to tumor cells with and/or healthy cells, likely by recognizing yet unknown cell-endogenous antigens (*9-11*).

MR1T cells use a polyclonal T-cell receptor (TCR) repertoire, can exhibit diverse T-helper-like functions (*10*) and are able to efficiently recognize various tumor cell types (*10*) and one clone showed broad tumor killing (*11*). These findings indicated MR1T cells as promising candidates for cancer immunotherapy (*12*). However, to enable the potential translation of MR1T cells into the clinic it is essential to understand the molecular targets they recognize in both physiological and pathological settings.

We hypothesized that the MR1T-cell antigens could be cell-endogenous metabolites, and focused our search on tumor cells, in which profound metabolic dysregulation occurs, that promote the generation and accumulation of unusual metabolites (*13, 14*). Using wide gene deletion technology to disrupt the metabolism of a model tumor target cell line we identify key cellular processes influencing MR1T-cell recognition. Next, we validated these results using single-gene manipulation and pharmacological approaches. Finally, we identified nucleobase-containing adducts as a class of MR1T-cell antigens and validated their stimulatory capacity using synthetic analogs. Our results suggest a role of MR1T cells as surveyors of cellular metabolic integrity as they recognize a novel class of self-antigens linked to tumor metabolic dysregulation.

## Results

### Knock-out of single metabolic pathway genes within target cells impacts MR1T-cell recognition

We devised a genome-wide CRISPR/Cas9-mediated gene disruption approach (*15*) to identify the cellular pathways involved in target cell susceptibility to MR1T-cell killing.

We hypothesized that inactivation of genes involved in antigen biosynthesis would decrease MR1T-cell-mediated killing by reducing the generation of antigenic compounds, while disruption of genes associated with antigen degradation would increase the accumulation of antigenic compounds, thereby increasing MR1T-cell susceptibility.

We transduced A375 melanoma tumor cells stably engineered with the MR1 and CAS9 genes (A375-MR1 cells) with a library of single guide RNAs (sgRNAs) covering the total human genome to generate targets for the cytotoxic MR1T cell clone TC5A87. After sequential rounds of T-cell killing, each followed by a recovery phase, surviving cells were analyzed by amplicon-sequencing for enriched or depleted sgRNAs compared to the same cells not subjected to killing. We removed genes essential for A375-MR1 cell growth (*16*) from the dataset to limit the impact of T-cell-independent processes. We defined significantly enriched or depleted sgRNAs (FDR>0.05) based on a log2 fold-change cut-off relative to the top and bottom 1% in the control (Figure 1A). Our approach identified 243 sgRNAs targeting 237 unique genes that were enriched, including MR1 and beta-2-microglobulin (B2M), as expected, which are both essential for MR1-dependent antigen presentation (Figure 1A). We also found enrichment of sgRNAs specific for the adhesion molecules CD58 (LFA-3) and ICAM-1, which bind the T-cell surface molecules CD2 and CD11a, respectively, and have a fundamental role in mediating target cell recognition and killing (*17*). Furthermore, we identified significant depletion of 5,331 sgRNAs targeting 4,705 unique genes (Figure 1A). A far greater number of depletions compared to enrichments was expected, as the number of genes required for tumor cell survival in general is likely to outweigh the number of genes that would allow specific escape from T-cell-mediated killing. Despite this fact, when we applied binomial enrichment analysis of gene-ontology (GO) terms (*18, 19*) annotated to significant hits, we revealed a large number of both enriched and depleted gene targets (n=127 and 650, respectively), associated with Metabolic Processes (Figure 1B, C), suggesting that the coordinated activity of multiple genes involved in target cell metabolism in general promoted MR1T-cell recognition. These hits showed enrichment in Nucleobase and Nucleic Acid Metabolic Processes, arguing for a potential role for these metabolic pathways in enhancing MR1T-cell stimulation.

**Figure 1.**
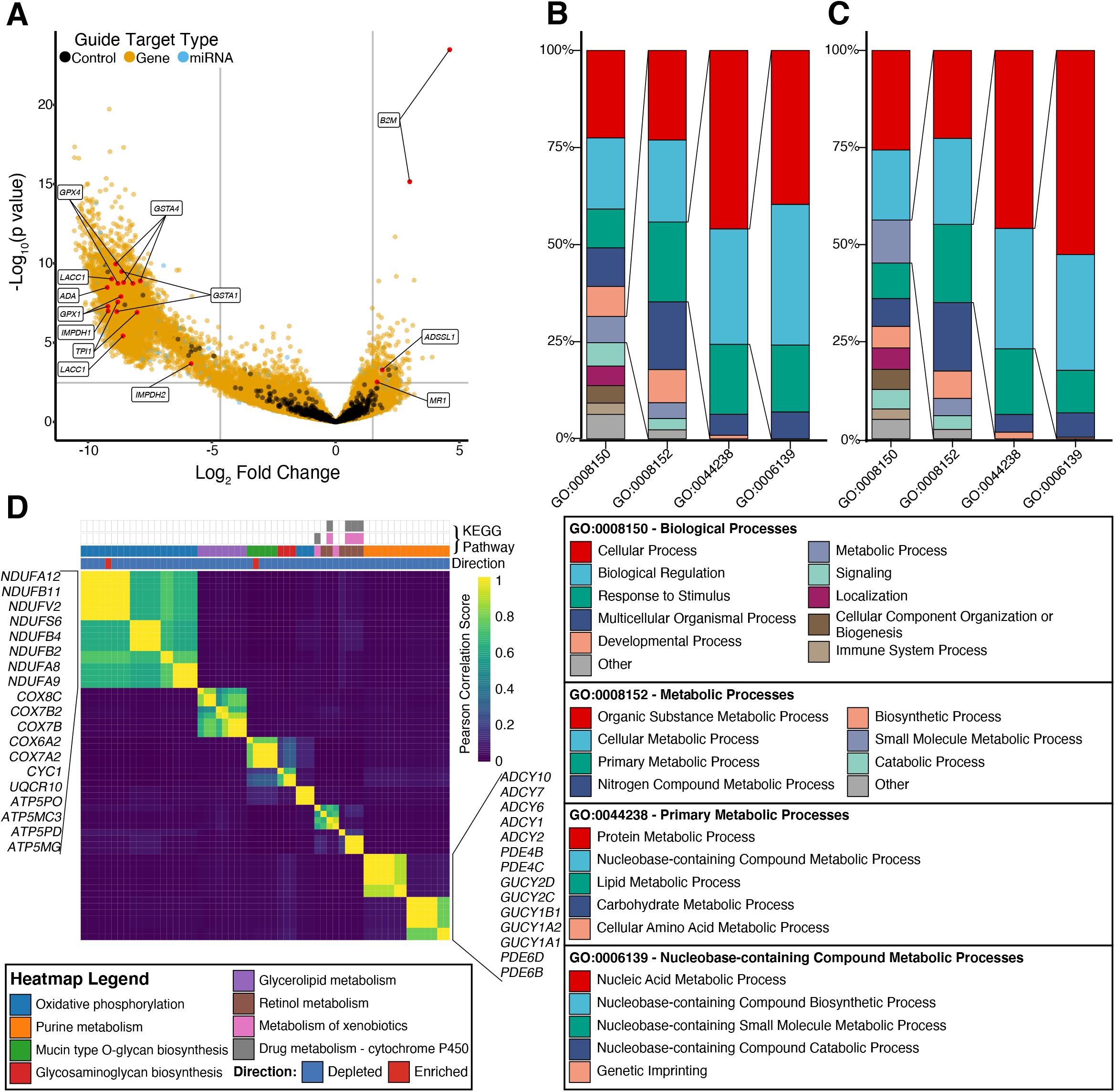
Identification and analysis of gene-targets influencing the efficiency of MR1T-cell mediated killing. (A) Significantly enriched and depleted guides were identified as having fold-change greater or lower than the top 1% of the negative control guides (black points), respectively (thresholds indicated by vertical grey lines) and FDR<0.05, here annotated as the highest *p*-value at which FDR is still <0.05 (horizontal grey line). (B and C) Significantly enriched (Binomial enrichment FDR<0.05) biological process GO-terms in enriched (B) and depleted (C) gene-targets. Each successive bar shows the significant child GO-terms. (D) Similarity of single-gene knockouts in the Recon3D metabolic perturbation model. See also Table S1, Data S1 and S2.

To explore the potential roles of the identified gene targets we used Recon3D, a genome-scale *in silico* model of human metabolism (*20*). We applied structural sensitivity analysis (*21, 22*) to predict the metabolic network response to single-gene knockouts (KOs). High Pearson correlations (score > 0.6) identified sets of two or more gene KOs predicted to have similar global effects on metabolic reactions (Figure 1D) and highlighted 125 genes among those identified by the CRISPR/Cas9 screening. Annotation of these genes with their KEGG pathways revealed that oxidative phosphorylation and purine metabolism were affected by single-gene knockouts of a number of different gene targets, leading to common metabolic responses (Figure 1D).

Taken together, our analyses of the CRISPR/Cas9 screen outlined the importance of several metabolic pathways within target cell for recognition by MR1T cells.

### Nucleobases presented by target cells can stimulate MR1T cells

We focused first on the purine pathway. The CRISPR/Cas9 screen identified several genes that might be involved in MR1T-cell antigen accumulation (Scheme S1, Data S1-S2). These genes were Adenosine Deaminase (*ADA*) that converts adenosine to inosine (*23*), Adenylosuccinate Synthase 1 (*ADSSL1*) necessary for the *de novo* production of adenosine monophosphate (AMP) from inosine monophosphate (IMP) (*24*), Laccase Domain Containing 1 (*LACC1*) that enables the purine nucleoside cycle (*25*), cGMP-specific 3’,5’-cyclic phosphodiesterase (*PDE5A*) that catalyzes the specific hydrolysis of cGMP to 5’-GMP (*26*), Aldehyde Dehydrogenase 16 Family Member A1 (*ALDH16A1)* and Hypoxanthine Phosphoribosyltransferase 1 (*HPRT1*), which both form a complex that generates purine nucleotides through the purine salvage pathway (*27*).

To validate the relevance of the selected genes, we measured the ability of individual KO cell lines to stimulate IFN-γ production by two different MR1T cell clones. *ADA*- and *LACC1-*deficient A375-MR1 cells induced significantly more IFN-γ production by both MR1T cell clones than did the parental A375-MR1 cells (Figures 2A, B, D, E), whereas the stimulatory ability of *ADSSL1*-deficient cells was significantly reduced, as predicted by the CRISPR/Cas9 screening (Figure 2C and 2F). The gene KOs did not impact on MR1 cell surface levels or 5-OP-RU presentation to MR1-restricted MAIT cells (Figure S1), thus excluding general alterations to the MR1 antigen-presentation pathway. These findings suggested that purines could be potential MR1T-cell antigens.

**Figure 2.**
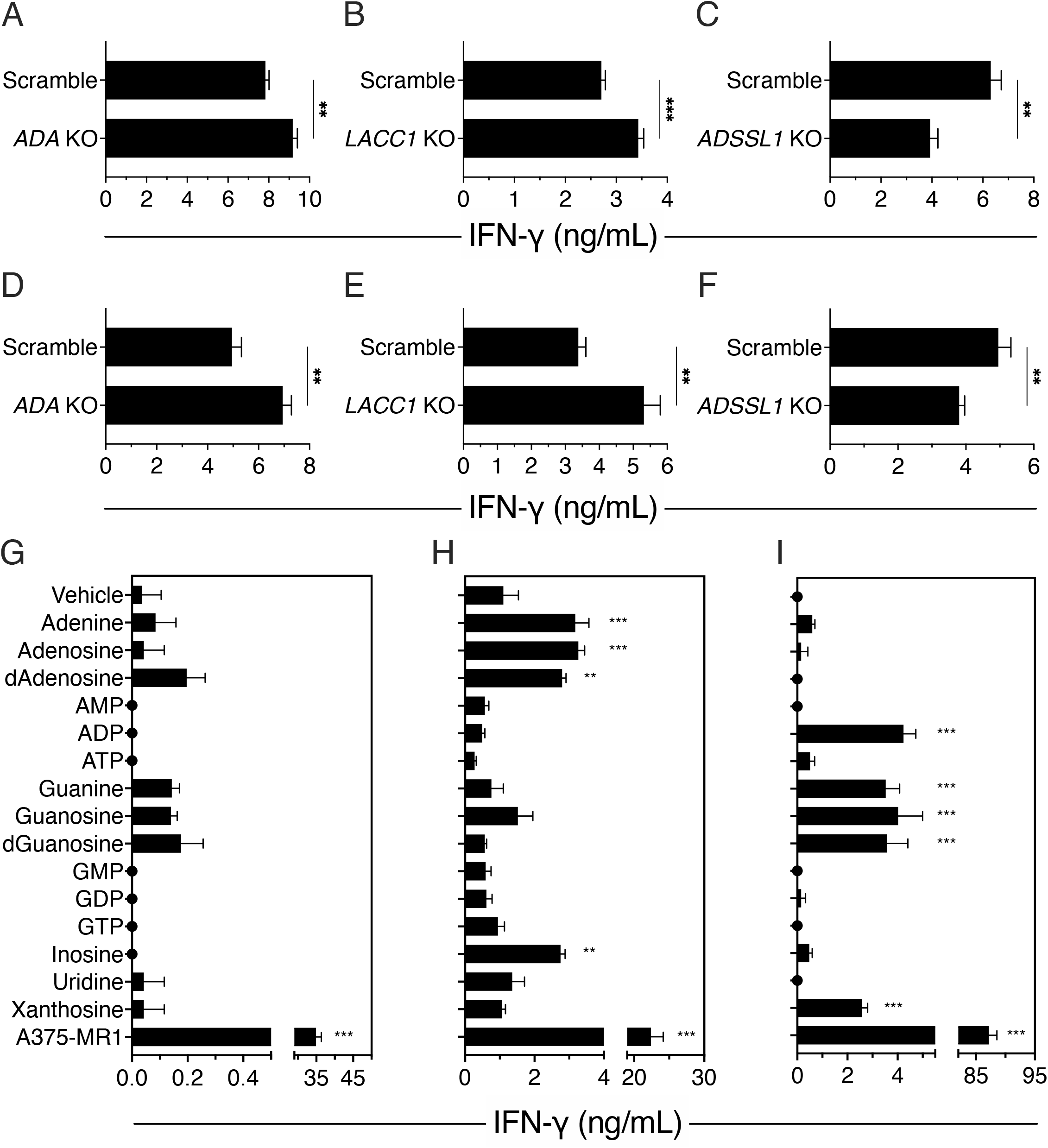
Purine metabolism is involved in MR1T antigen accumulation. (A-F) MR1T clones TC5A87 (A-C) and DGB129 (D-F) reactivity against A375-MR1 cells transduced with sgRNAs targeting ADA (A and D), LACC1 (B and E), ADSSL1 (C and F) or scrambled sgRNA control (A-F). (G-I) Activation of MR1T clones TC5A87 (G), DGB129 (H), and MCA3C3 (I) by THP-1 cells pre-incubated with 250 µM of the indicated molecules or vehicle. A375-MR1 cell stimulation is shown as positive control. IFN-γ released is the mean ± SD of triplicate cultures. One representative experiment of at least three independent replicates is shown in each panel. **p ≤0.01 and ***p ≤0.001 compared to matching control (A-F, Unpaired t-test) or compared to vehicle (G-I, One-way Anova with Dunnett’s multiple comparison). See also Figures S1, S2 and Scheme S1.

Next, we incubated human acute monocytic leukemia THP-1 cells with synthetic nucleotides, nucleosides or nucleobases before adding MR1T-cell clones and measured IFN-γ production. THP-1 cells were selected as targets because they constitutively express low surface levels of MR1 and induce some spontaneous MR1T-cell stimulation, demonstrating their ability to appropriately process and present MR1T-cell antigens. A375-MR1 cells expressing very high surface levels of MR1 were also included as a positive control to stimulate MR1T-cell cytokine production. Three distinct MR1T-cell clones differently reacted to tested compounds: TC5A87 did not show significant reactivity (Figure 2G); DGB129 reacted to adenine, adenosine, deoxyadenosine (dAdo) and inosine (Figure 2H); and MCA3C3 was activated by ADP, guanine, guanosine, deoxyguanosine and xanthosine (Figure 2I). THP-1 cells incubated with the synthetic compounds did not stimulate MAIT cells (Figure S2A). Interestingly, the stimulatory effect of these compounds was observed at high doses, leading us to hypothesize that they could be intermediate precursors of antigens.

### Methylglyoxal and purine metabolism pathways within target cells cooperate for MR1T-cell stimulation

To understand whether additional metabolic pathways might be involved in MR1T cell reactivity to nucleobases/nucleosides we interrogated our whole-genome gene disruption screening data. We observed that some of the significantly depleted sgRNAs were related to genes involved in glycolysis, such as triosephosphate isomerase 1 (*TPI1*) and genes involved in methylglyoxal (MG) degradation, including Glyoxalase 1 (*GLO1)*, and Glyoxalase Domain Containing 4 (*GLOD4*). *TPI1* is responsible for the enzymatic conversion of dihydroxyacetone phosphate (DHAP) into glyceraldehyde 3-phosphate (G3P) — a reaction that can otherwise occur spontaneously with the generation of MG (*28*) (Figure 3A). Conversely, *GLO1*-deficient cells are impaired in MG (a highly reactive carbonyl) degradation, which therefore accumulates (*29*). As MG forms adducts with several nucleobases (*30*), these data suggest a potential involvement of MG in generating MR1T-cell antigens.

**Figure 3.**
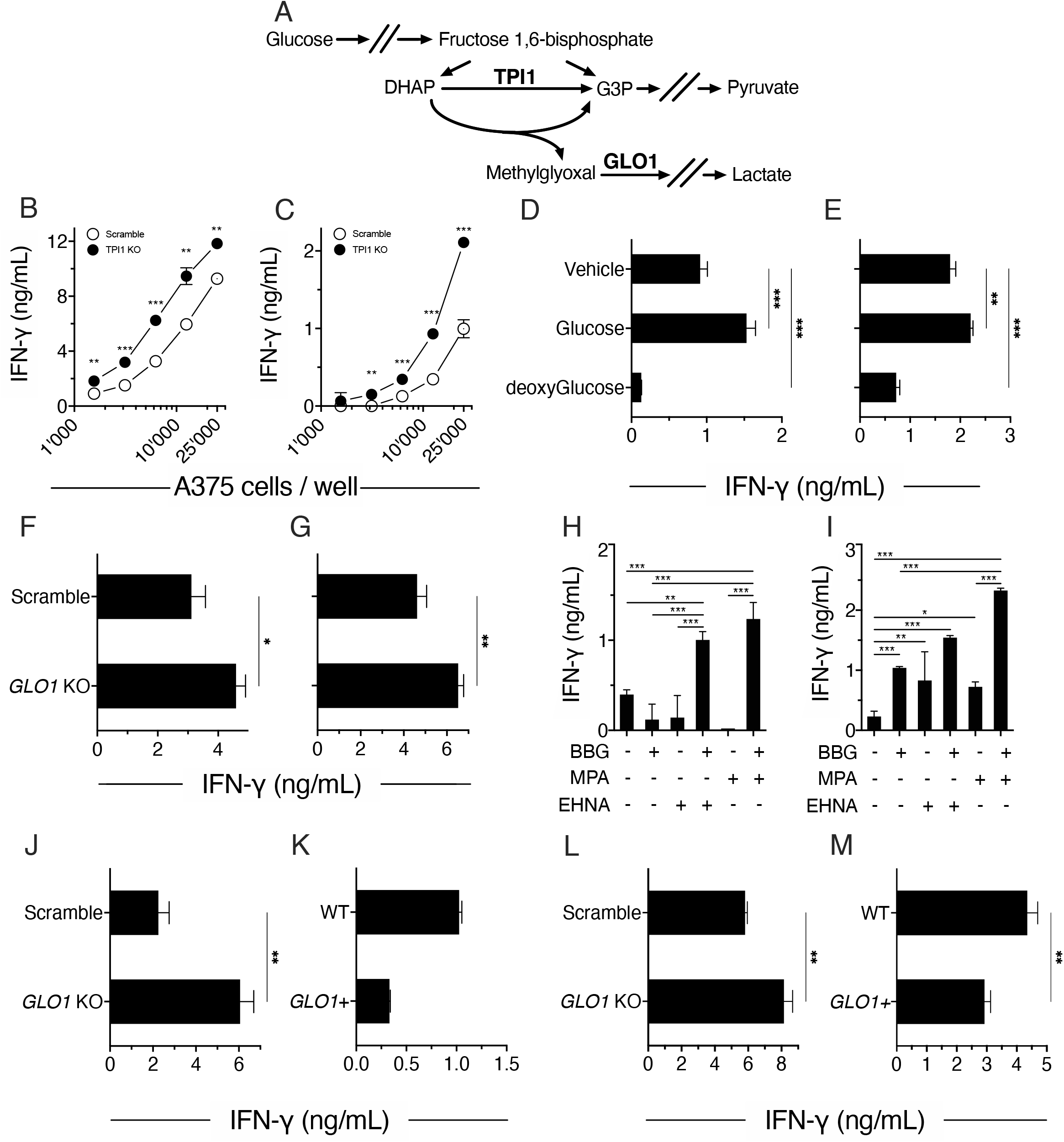
Glycolysis and methylglyoxal contribute to MR1T cell stimulation. (A) Schematic representation of methylglyoxal generation. Dihydroxyacetone phosphate (DHAP), Glyceraldehyde 3-phosphate (G3P), Triosephosphate isomerase (TPI1), and Glyoxalase 1 (GLO1). (B and C) Stimulation of MR1T cell clone TC5A87 (B) and DGB129 (C) with A375-MR1 cells transduced with sgRNAs targeting TPI1 (•) or scrambled control (○). (D and E) Stimulation of MR1T cell clone TC5A87 (D) and DGB129 (E) in response to fixed A375-MR1 cells incubated with D-(+)-Glucose (15 mM) or 2-deoxy-D-Glucose (15 mM) before fixation. (F and G) Stimulation of MR1T cell clone TC5A87 (E) and DGB129 (F) with A375-MR1 cells transduced with sgRNAs against GLO1 or scrambled sgRNAs controls. (H and I) Stimulation of MR1T cell clone TC5A87 (H) and DGB129 (I) with THP-1 cells pre-treated with EHNA, MPA and BBG, alone or in combination. (J-M) MR1T clone DGB129 activation in response to THP-1 cells unmodified (WT), overexpressing *GLO1* (*GLO1*^+^), transduced with *GLO1*-targeting (*GLO1* ko) or scrambled control (Scramble) sgRNAs, in the presence of methylglyoxal (50 µM, J and K) or deoxyadenosine (1 mM, L and M). IFN-γ released is the mean ± SD of triplicate cultures, except for K which is in duplicate. The data shown are representative of at least three independent experiments. (B-M) *p<0.05 **p ≤0.01 and ***p ≤0.001. (B and C) Multiple t-test, (D and E) one-way Anova with Dunnett’s multiple comparison, (F, G and J-M) unpaired t-test, (H and I) one-way Anova with Tukey’s multiple comparison. See also related Figures S1 and S2.

We then dissected the possible roles of glycolysis and MG degradation in MR1T stimulation by generating single gene KO cell lines. Loss of triosephosphate isomerase protein in A375-MR1 cells significantly increased IFN-γ production by both MR1T-cell clones (Figure 3B, C). Furthermore, A375-MR1 cells pulsed with glucose and then fixed showed increased stimulatory capacity (Figure 3D, E); this effect was abolished when the same cells were incubated with deoxyglucose (Figure 3D, E), which does not enter the glycolytic pathway and so does not generate MG (*31*). We also saw that GLO1-deficient A375-MR1 cells showed increased stimulatory capacity (Figure 3F, G). Altogether these results suggest that MG accumulation in target cells is important for the stimulation of MR1T-cell clones.

To investigate the potential synergism between MG and purine metabolic pathways in MR1T-cell stimulatory capacity of tumor cells, we explored the effects of individual and combined pharmacologic inhibition of key enzymes in these pathways (Scheme S1). We used S-*p*-bromobenzylglutathione (BBG) to inhibit GLO1 (*32, 33*); mycophenolic acid (MPA) to inhibit inosine monophosphate dehydrogenases (IMPDH1, and IMPDH2) (*34*), leading to IMP accumulation; and erythro-9-(2-Hydroxy-3-nonyl) adenine hydrochloride (EHNA) to inhibit ADA (*35*) and phosphodiesterase 2 (PDE2) (*36*), inducing adenosine, dAdo and cGMP accumulation (*35*). To sensitively detect the effects of the inhibitors, we again used THP-1 cells as target cells: we found that BBG in combination with each of the other two drugs significantly enhanced the IFN-γ release of both MR1T cell clones to THP-1 cells (Figure 3H, I). The DGB129 clone was more sensitive to these treatments and also reacted to THP-1 cells treated with EHNA, BBG, or MPA alone (Figure 3I).

Finally, we tested the IFN-γ response of MR1T cells to GLO1-modified THP-1 cells and MG or dAdo. We found that MG treatment significantly increased stimulation by GLO1-deficient compared to wild-type THP-1 cells (Figure 3J). Conversely, MG failed to induce stimulation when administered to GLO1-overexpressing cells (Figure 3K). Similarly, MR1T reactivity to dAdo was increased using GLO1-deficient THP-1 cells as antigen-presenting cells (APCs) (Figure 3L) and decreased using GLO1-overexpressing THP-1 cells (Figure 3M). Together, these findings suggest that nucleosides/nucleobases and MG cooperate in generating potential MR1T-cell antigens.

### Oxidative stress-related carbonyl species accumulating within target cells contribute to MR1T cell stimulation

In addition to the purine pathway, our model-based analysis (Figure 1D) highlighted genes related to oxidative phosphorylation, whose protein products participate in ATP generation within mitochondria and whose alteration promotes accumulation of reactive oxygen species (ROS) (*37, 38*). Alongside, the analysis pointed towards the relevance of the H^+^ transporter subunits ATP6V1C2, TCIRG1 and ATP6V0D2 involved in coupling proton transport and ATP hydrolysis and thus contributing to maintaining the organelle physiological milieu in the cell, including mitochondria (*39*). Therefore, we next investigated the roles of ROS in MR1T-cell stimulation by tumor cells.

First, we focused on genes involved in oxidative phosphorylation. Our initial MR1T-cell killing screen uncovered a significant depletion of sgRNAs specific for *GSTM1, GSTA4, GSTA1, GSTM5, GSTA2, GSTM3*, and *GSTO1* (Data S2). The products of these genes are involved in the detoxification of electrophilic compounds and ROS by their conjugation to glutathione (GSH), a ROS scavenger (*40, 41*). We therefore hypothesized that in the absence of GSTs, and upon accumulation of ROS and electrophilic molecules (*42*) (Scheme S2), tumor cells may accumulate MR1T-cell stimulatory compounds. Accordingly, we tested the effects of paclitaxel and doxorubicin, two drugs that induce cellular accumulation of O_2-_ and H_2_O_2_ (*43-45*). Both drugs significantly increased ROS accumulation (Figure S3) and promoted activation of all three MR1T-cell clones when incubated with THP-1 cells and nucleoside compounds (Figure 4A-C), mirroring the additive effects observed when purine-modifying drugs and carbonyl-degradation inhibitors were combined (Figure 3H, I).

**Figure 4.**
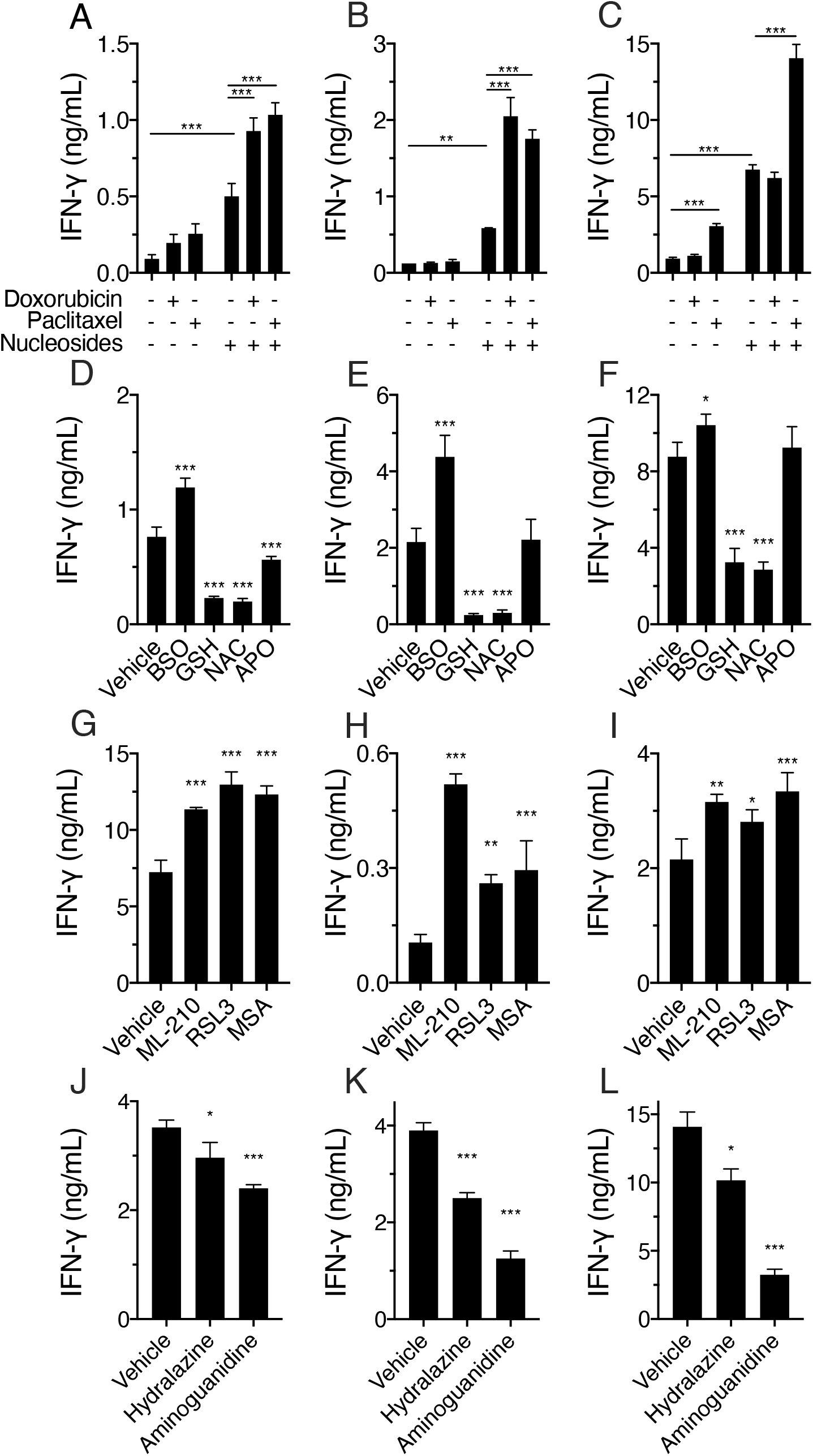
Endogenous aldehyde alterations contribute to MR1T cell stimulation. (A-C) Stimulation of MR1T cell clone TC5A87 (A), DGB129 (B) and MCA3C3 (C) with THP-1 cells pre-treated with Doxorubicin or Paclitaxel in the absence or presence of nucleosides (dAdenosine and Guanosine). (D-F) Stimulation of MR1T cell clone TC5A87 (D), MCA2B1 (E) MCA3C3 (F) with fixed A375-MR1 cells treated with BSO, GSH, NAC, APO or vehicle. (G-I) Stimulation of MR1T cell clone TC5A87 (G), DGB129 (H) MCA2B1 (I) with fixed A375-MR1 cells treated with ML-210, RSL3 or MSA. (J-L) Stimulation of MR1T cell clone TC5A87 (J), DGB129 (K) and MCA2B1 (L) with fixed A375-MR1 cells treated with Hydralazine or Aminoguanidine. IFN-γ release is the mean ± SD of triplicate cultures. The data shown are representative of at least three independent experiments. *p<0.05, **p ≤0.01 and ***p ≤0.001. (A-C) two-way Anova with Tukey’s multiple comparison, (D-L) one-way Anova with Dunnett’s multiple comparison. See also Figures S2, S3 and Scheme S2.

Next, we treated A375-MR1 with apocynin, an O_2-_ scavenger and NADPH oxidase inhibitor (*46*), or with GSH or N-acetylcysteine (NAC), which prevent H_2_O_2_ accumulation (*43*) before fixation and incubation with the three MR1T cell clones. We found that A375-MR1 cells treated with GSH or NAC stimulated significantly less IFN-γ production from MR1T cells, whereas apocynin was less active and inhibited one MR1T clone only (Figure 4D-F). We also treated A375-MR1 cells with buthionine sulfoximine (BSO), an inhibitor of GSH synthase (*47*), and a significant increase was observed in the stimulation of all the tested MR1T clones (Figure 4D-F). Together, these data show that ROS accumulation contributes to MR1T cell stimulation, with maximal effects in the presence of nucleobases increase.

Peroxide accumulates in many tumor types and is involved in various signal transduction pathways and cell fate decisions (*48*). Peroxide is also necessary for lipid peroxidation, a pathway that generates malondialdehyde (MDA) and 4-OH-nonenal (4-HNE), two highly reactive carbonyls (*49, 50*). Both compounds form stable adducts with proteins, lipids and nucleobases and accumulate within tumor cells (*51-54*). Alongside our finding that inhibiting ROS accumulation impedes target cell stimulation of MR1T cells (Figure 4D-F), we inferred a role for lipid peroxidation from the results of our CRISPR/Cas9 screen, which also showed significant depletion of the glutathione peroxidase 4 (GPX4) and glutathione peroxidase 1 (GPX1) sgRNAs (Figure 1 and Data S2). While GPX1 protein catalyzes the reduction of organic hydroperoxides and H_2_O_2_ by glutathione, GPX4 has a high preference for lipid hydroperoxides (Scheme S2) and protects cells against membrane lipid peroxidation and death (*55*). Accordingly, when we pre-treated A375-MR1 cells with the selective GPX1 inhibitor mercaptosuccinic acid (MSA), or with two GPX4 inhibitors RSL3 and ML-210, they showed significantly increased MR1T-cell stimulatory activity (Figure 4G-I). None of these compounds induced activation MAIT-cell, except paclitaxel that in presence of nucleosides could induce a little but significant stimulation of one MAIT clone (Figure S2C, D). Taken together, these findings indicate that peroxides and lipid peroxidation contribute to MR1T-cell stimulation by target cells.

To further assess the carbonyl involvement in the generation of MR1T antigens, we tested the capacity of carbonyl scavengers to prevent MR1T-cell activation. A375-MR1 cells were incubated with aminoguanidine and hydralazine, which show preferential scavenging activity for different carbonyls (*56*), then fixed and washed before the addition of MR1T cell clones. Both scavengers significantly inhibited IFN-γ production by MR1T cell clones (Figure 4J-L) and had no effect on MAIT cell activation (Figure S2E).

Collectively, our data suggest that multiple oxidative stress-related reactive carbonyl species accumulating in cells following metabolic alterations combine with nucleobases to generate MR1-presented antigens that stimulate MR1T cells. The biochemical condensation of carbonyls with nucleobases physiologically occurs in cells, and in tumors leads to the accumulation of adducts responsible of DNA damage (*57*). Carbonyl adducts of nucleobases are present within tumor cells and also accumulate in pre-cancer lesions (*58-61*). Thereby, we tested whether nucleobase-carbonyl adducts stimulate MR1T cells.

### Nucleobase carbonyl adducts stimulate MR1T cells

Three nucleobase-carbonyl adducts were tested for their activation capacity. These compounds were selected because: i) they have been described to be present within cells (*54, 62-64*), ii) represent adducts of three different nucleobases, and iii) they are formed by condensation of the nucleobase with MDA. This carbonyl is a product of cellular oxidative stress and lipid peroxidation (*65*), is abundant within tumor cells (*53*) and readily reacts with different nucleobases (*66*). The three compounds were: *N*^6^-(3-oxo-1-propenyl)-2’-deoxyadenosine (OPdA) (*67*), pyrimido[1,2-*α*]purin-10(3*H*)-one (M_1_G) (*66, 68*), and the pyrimidine adduct, *N*^4^-(3-oxo-1-propenyl)-2’-deoxycytidine (OPdC) (*69*). The NMR and MS spectra are reported in Figure S4, and S5.

To test whether these carbonyl nucleobase adducts might behave as MR1T antigens, we investigated compound-induced MR1 upregulation and MR1T cell activation assays using synthetic analogs presented by live APCs or by recombinant soluble MR1.

All three compounds induced upregulation of MR1 (Figure 5A-C, middle panels). Their antigenicity was investigated using as responders a panel of MR1T cell clones. Each compound was stimulatory when presented by live APCs (Figure 5A-C, right panels), thus showing their antigenicity. In all cases, MR1T-cell activation by adduct-loaded THP-1 cells was fully inhibited by the addition of blocking anti-MR1 monoclonal antibodies (mAbs) (Figure 5A-C, right panels), confirming MR1 restriction of adduct recognition.

**Figure 5.**
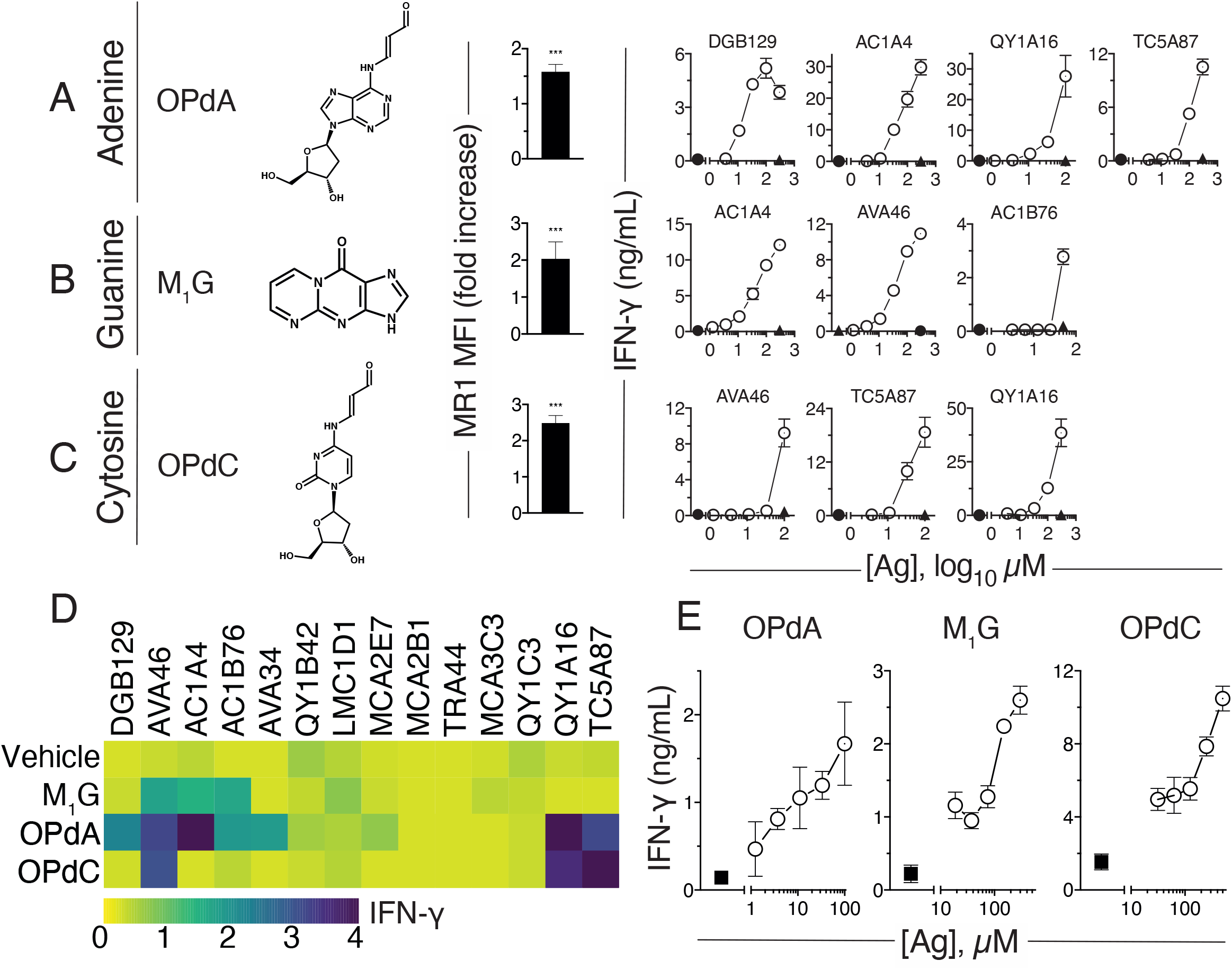
Synthetic MDA nucleoside adducts stimulate MR1T cells. (A-C) (Left), structure of three synthetic adducts OPdA (A), M_1_G (B), and OPdC (C). (Middle), upregulation of MR1 surface expression on THP-1 cells following 6 h incubation with each adduct. Fold increase of median fluorescence intensity (MFI) ± SD compared to cells treated with vehicle is shown. (Right), IFN-γ release response of indicated MR1T cell clones co-cultured with THP-1 cells in the presence (○) or absence of adducts (•). Blocking of T-cell reactivity is shown for the highest antigen (Ag) dose using anti-MR1 mAbs (▴). (D) Stimulation of fourteen MR1T cell clones in the presence of THP-1 cells and OPdA, M1G, OPdC, or vehicle. Heat-map reports the square root of mean IFN-γ concentration. Antigen response was compared to vehicle and evaluated by multiple t-test (p<0.05). (E) Response of MR1T clones QY1A16, AC1A4 and TC5A87, stimulated by plate-bound soluble MR1 loaded (○) with OPdA, M_1_G and OPdC, respectively, or not loaded (▪). The data shown are representative of at least three independent experiments. (A-C) Mean ± SD, n≥3. (D-E) Mean ± SD, n≥2. See also related Figures S2, S3, S4 and S5.

We further confirmed the stimulatory capacity of these nucleobase adducts by testing additional MR1T-cell clones expressing different TCRs: of the fourteen randomly selected MR1T clones tested, three reacted to M_1_G, seven to OPdA and three to OPdC (Figure 5D). Of note, THP-1 cells pulsed with the synthetic compound did not stimulate the MAIT-cell clone MRC25 (Figure S2F). These findings indicated that the same MR1T TCRs crossreact with different nucleobase adducts. However, other TCRs did not recognize the tested antigens, indicating their specificity for other antigens.

In additional experiments, we used plastic-bound recombinant MR1 molecules loaded with synthetic antigens. These experiments showed that all tested compounds were stimulatory, and MR1T cell stimulation was blocked by addition of anti-MR1 mAbs (Figure 5E). Thus, carbonyl-nucleobase adducts bind MR1 without further processing by APC and stimulate MR1T-cells.

In conclusion, we show that a group of MR1T cells recognize compounds containing intact carbonyl adducts of nucleobases presented on MR1 molecules. Some MR1T cell clones preferentially recognize one or another adduct; others show a degree of cross-reactivity. This promiscuous recognition may promote broad reactivity of MR1T cells.

## Discussion

In this study, we have identified nucleobase adduct-containing metabolites as a class of self-antigens capable of stimulating human T lymphocytes. Previous studies showed that MR1T cells react to unique compounds fractionated from tumor cells, suggesting distinct antigen specificity (*10*). Here, we confirm those data and extend them to show that structurally-diverse nucleobase adduct-containing compounds bind MR1 and stimulate individual MR1T cells. Both purines and pyrimidines can form antigenic adducts and different carbonyls participate in their generation, confirming that MR1 is a molecule with versatile antigen-binding capacity.

Carbonyls accumulate as a consequence of different metabolic alterations in glycolysis and lipid peroxidation, during the metabolism of biogenic amines, vitamins and steroids, as well as upon biotransformation of environmental agents and drugs (*54, 70, 71*). How many carbonyls are involved warrants further investigation: this is an important question as the number and diversity of carbonyl species participating in the generation of MR1-presented nucleobase adducts may determine the size and variety of the antigen repertoire recognized by MR1T cells responding to this class of molecules. It is of note that some MR1T clones did not react to any of the three antigens tested here, whereas still responded to A375-MR1 cells, suggesting the MR1T cell antigen repertoire can be quite large and heterogeneous, probably reflecting the variety of nucleobase adducts accumulating within tumor cells (*57*).

While carbonyl accumulation is important, it is not sufficient to stimulate MR1T cells, as concomitant availability of free purines and pyrimidines must occur within the target cells. The structures of modified nucleobases are suited to MR1 binding and resemble those of other MR1 ligands, as they are composed of differently modified heterocyclic compounds (*72*). Future studies will certainly shed light on the repertoire size of the MR1-binding molecules that stimulate MR1T cells.

Additional important players in the generation of immunogenic nucleobase adducts are ROS, that promote lipid peroxidation leading to carbonyl generation (*73*), thus also potentially exerting indirect effects on accumulation of MR1T cell antigens. Indeed, treatment with ROS-inducing drugs enhanced MR1T-target cell stimulatory capacity, which in turn was dampened by the addition of ROS scavengers. Thus, a class of MR1T-cell antigens are derived by combined alterations of distinct metabolic pathways, co-leading to the accumulation and condensation of nucleobases and carbonyls.

MR1T-cell recognition of nucleobase adduct-containing metabolites raises the question of the physiological role of these cells. It is conceivable to attribute them a potential role in surveying cells abnormally undergoing metabolic reprogramming and, therefore, generating and/or accumulating compounds responsible for nucleobase alterations (*74*). The ubiquitous expression of MR1 might be instrumental to this function of cellular metabolic integrity control. Together these properties of MR1T cells make them attractive targets for immunotherapeutic use in diseases associated to multiple metabolic dysregulations, concomitantly affecting nucleobase homeostasis, oxidative status and carbonyl accumulation, such as cancer.

An important, still unanswered question in the field has been the basis of tumor cell recognition by MR1T cells. Transformed cells frequently become altered in several pathways involved in nucleobase generation and sustaining cell proliferation, including alterations in glucose and glutamine uptake, and high cellular demand for reduced nitrogen (*75*). Indeed, many tumors increase the transcription of key genes involved in the *de novo* purine synthesis (*76*). Furthermore, tumor cells are prone to DNA damage through the generation of nucleic acid adducts formed upon DNA oxidation and interaction with end products of lipid peroxidation (*77*). The combined presence of these alterations may promote accumulation of antigenic nucleobase adducts in tumor cells. Understanding the relative contribution of these metabolic alterations to MR1T antigen accumulation may pave the way to HLA-unrestricted and broadly tumor-reactive MR1T cell-based immunotherapies for cancer.

## Acknowledgements

We thank Verena Schaefer, and Florine Winter for contributing to experiments, Feng Zhang for Addgene Human GeCKO v2 knockout library and plasmids 52962-3. We wish to thank Lucy Robinson and Jessica Tamanini of Insight Editing London for assistance in preparing the manuscript. Work in De Libero’s group was supported by grants of Swiss National Foundation (310030-173240 and 310030B-192828), Swiss Cancer Research foundation (KFS-4707-02-2019), Cancer League beider Basel (KLbB-4779-02-2019), D-BSSE ETH Zürich (PMB-02-17) and by the University and University Hospital of Basel.

## Author Contributions

Conceptualization and Methodology, L.M. and G.D.L.; Investigation, A.V., A.C., J.S. Q.Y., D.J., A.B., C.D.G., M.P. and M.L.; Formal analysis, O.Ø., J.St.; Resources, G.D.L.; Data curation, A.V., A.C., J.S., D.H.; Writing original draft, A.V., A.C., J.S., G.D.L.; Writing-Review and Editing, M.L., L.M. and G.D.L.; Visualization, A.V., A.C., Q.Y., J.S., D.J.; Supervision, L.M and G.D.L.; Funding Acquisition, G.D.L. All authors approved the manuscript.

## Supplemental Information

Supplemental information includes 5 figures, 2 tables, 2 schemes and 2 data files.

Figure S1. Genetically-engineered cell lines characterization, Related to Figures 2 and 3.

Figure S2. Stimulation of MAIT clone MRC25 with nucleobases, inhibitory drugs and synthetic antigens, Related to Figures 2-5.

Figure S3. ROS accumulation in APCs and MR1 expression in tumor cell lines, Related to Figure 4.

Figure S4. Mass-Spectrometric and NMR analyses of synthetic OPdC and OPdA, Related to Figure 5.

Figure S5. Mass-Spectrometric and NMR analyses of synthetic M1G, Related to Figure 5.

Table S1. KEGG pathways related to statistically significant enriched or depleted sgRNAs, Related to Figure 1, Data S1 and S2.

Table S2. sgRNA target sequences used for knock-out generation, Related to Figures 2 and 3.

Scheme S1. sgRNA hits identified in purine metabolism pathway, Related to Figures 1-3 and Data S1 and S2.

Scheme S2. ROS generation and scavenging, Related to Figures 1, 4 and Data S1 and S2.

Data S1. Significantly enriched sgRNAs classification according to KEGG pathway, Related to Figure 1 and Table S1.

Data S2. Significantly depleted sgRNAs classification according to KEGG pathway, Related to Figure 1 and Table S1.

## EXPERIMENTAL MODEL AND SUBJECT DETAILS

### Human blood samples

Blood specimens for T-cell cloning, FACS analysis and antigen-presentation assays were obtained from the University Hospital Basel. The study was approved by the local ethical review board (EKNZ, Ethics Committee North-West & Central Switzerland, EKNZ 2017-01888), and all healthy donors consented in writing to the analysis of their samples.

## METHOD DETAILS

### Cell culture

A375-MR1 and THP-1-MR1 cells were generated as previously described (*10*). Cell lines were cultured in RPMI-1640 supplemented with 10% FCS, 2 mM L-glutamine, 1 mM sodium pyruvate, 1x MEM NEAA and 50 µg/ml kanamycin (all from Bioconcept). All human T-cell clones were maintained in culture as previously described (*78*). A representative MAIT clone (MRC25) generated from blood of a healthy donor was previously characterized (*78*). Cells were free from Mycoplasma as evaluated by PCR analysis on DNA samples. When possible, cells were authenticated by staining with mAb for specific cell surface markers.

Lentiviral transductions were carried out as previously described (*10*). Transduced cells were selected by FACS based on the expression of EGFP or mCherry reporters, or by puromycin resistance.

### Human knockout library screening

A375-MR1-Cas9 cells generated using the previously described cell line (*10*) and Lenti Cas9-Blast plasmid (Addgene, Cat#52962) were transduced at 0.3 MOI by both part A and B of the pooled Human GeCKO v2 CRISPR library (Addgene, Cat#1000000049), and subsequently selected using 2 µg/mL puromycin (Calbiochem, Cat#540411) for 96 h. Eight biological replicates of the resulting APCs, each with 64-fold representation of each guide within the library, underwent four consecutive rounds of killing by TC5A87 cells at a E:T of 2:1. The surviving cells were expanded for 24 h and DNA was extracted using a NucleoSpin Tissue kit (Macherey-Nagel, Cat#740952). An additional eight biological replicates were similarly prepared but did not undergo killing by TC5A87, as to serve as controls. Illumina libraries were prepared as previously described (*79*). Briefly, primers JScrispr1 (5’ TCGTCGGCAGCGTCAGATGTGTATAAGAGACAGNNNNNNNNGCTTTATATATCTTGTGGAAAGGACGAAACACC) and JScrispr3 (5’ GTCTCGTGGGCTCGGAGATGTGTATAAGAGACAGCTGCCATTTGTCTCAAGATCTAGTTACGCCAAG) were used to amplify genomic sgRNA from the extracted gDNA and attach common Illumina primer handles for attaching sequencing library indexes. The former primer also inserts an 8-nt degenerate sequence immediately downstream of the Illumina read 1 start site, decreasing issues of sequencing low-complexity libraries. Each replicate was barcoded by a unique pair of Nextera indexes (Illumina, Cat#15055290) in a second step PCR performed, as described in the Nextera DNA library preparation protocol (Illumina). A high-fidelity Advantage HF2 PCR kit (Takara, Cat#639123) was used in each of the PCR steps involved in preparing the sequencing libraries. Libraries were quantified using a BioAnalyser high sensitivity DNA kit (Agilent, Cat#5067-4626) and a Qubit high-sensitivity dsDNA kit (ThermoFisher, Cat#Q32851) and pooled to form an equimolar sequencing library that was denatured and diluted to 1.2 pM with a 20% PhiX v3 control library (Illumina, Cat#FC-110-3001), as described in the Illumina Denature and Dilution protocol (Illumina) before sequencing on a NextSeq500 using a High-output 150-cycle v2 kit (Illumina, discontinued product). Both sets of sequencing libraries were sequenced using a dual-indexed single-end protocol (131 cycles on read 1, 8 cycles on each barcode) to a depth of 25 million reads per replicate, ensuring that guides depleted after T-cell mediated killing may be detected.

### CRISPR/Cas9-mediated gene disruption

Results obtained in the screening were confirmed by knock-out of selected genes in A375-MR1-Cas9 cells transduced with sgRNAs different from the ones present in the library (Table S2). After lentiviral transduction and selection, A375-MR1-Cas9 cells were maintained for a limited number of passages and used in activation assays as bulk populations. The expression levels of target proteins were assessed by western blotting (Figure S1) with antibodies against ADA (clone D1P4Y, Cell Signaling Technology, Cat#65184), ADSSL1 (clone 2D12, Sigma-Aldrich, Cat#SAB1401979), Glyoxalase 1 (clone 266, Novus Biologicals, Cat#NBP2-43618), LACC1 (clone E-12, Santa Cruz Biotechnologies, Cat#sc-376231) TPI1 (Proteintech, Cat#13937-1-AP), Actin (clone C-2, Santa Cruz Biotechnologies, Cat#sc-8432) or α-Tubulin (clone B-5-1-2, Sigma-Aldrich, Cat#T5168). Primary antibodies were revealed with IRDye® 680 Goat anti-Mouse IgG or IRDye® 800CW Goat anti-Rabbit IgG (LI-COR, Cat#926-32220 and Cat#926-32211, respectively) or HRP Goat anti-Mouse IgG (H+L) (ThermoFisher, Cat#A16072) and WesternBright Sirius (Advansta, Cat#K12043). MR1 surface expression was evaluated by flow cytometry with APC-labeled mouse anti-MR1 mAbs 26.5 (Biolegend, Cat#361108) and APC-labeled mouse IgG2a (clone MOPC-173 Biolegend, Cat#400220) as isotype control (Figure S1). The antigen presentation ability of different cell lines was tested by stimulation of MAIT clone MRC25 after pulsing APCs for 2 h at 37°C with indicated concentrations of 5-OP-RU freshly-prepared with 5-A-RU (Toronto Research Chemicals, Cat#A629245) and Methylglyoxal (MEG, Sigma-Aldrich, Cat#M0252) (Figure S1).

### Preparation and purification of synthetic antigens

Synthetic compounds were synthesized and subsequently purified before use with cells.

#### Synthesis of Pyrimido[1,2-a]purin-10(3H)-one (M_1_G, CAS 103408-45-3)

M_1_G was synthesized as previously described (*80*) and (*81*) with some modifications. 1,1,3,3-Tetraethoxypropane (1.4 g, 6.25 mmol, 5.0 eq.) in aq. HCl (25 mL, 1 M) was stirred at 40°C for 1 h. Subsequently, a solution of guanine (188.9 mg, 1.25 mmol, 1.0 eq., Sigma-Aldrich, Cat#G11950) in aq. HCl (25 mL, 1 M) was slowly added. The mixture was stirred at 40 °C for 1 h and then kept at 4 °C for 16 h. The precipitate was washed three times with absolute ethanol at 2000 x *g* for 10 min. The raw M_1_G was extracted three times from the precipitate with 65°C water. The combined extracts were filtered with 0.22 µm filter. The mixture was adjusted to pH 7.0 with aq. NaOH (1 M).

M_1_G HPLC purification was performed as for M_3_ADE purification using the same type of HPLC column and mobile phases. Separation was performed with a flow rate of 6 mL/min and a linear gradient of 0-20% B from 0 to 40 min, 20-100% B from 40 to 41 min, 100% B from 41 to 46 min, 100-0% B from 46 to 47 min and 0% B until 55 min. M_1_G yield was 12.5 mg (66.8 μmol, 5.3%). Biologically active HPLC peaks were collected for mass spectrometric and NMR analyses.

^**1**^**H-NMR** (600 MHz, D_2_O, *δ*/ppm): 9.31 (d, ^*3*^*J*_*H13-H12*_ = 7.2 Hz, 1H, **H**_**13**_), 8.97 (dd, ^*3*^*J*_*H11-H12*_ = 4.1 Hz, ^*4*^*J*_*H11-H13*_ = 2.0 Hz, 1H, **H**_**11**_), 8.22 (s, 1H, **H**_**8**_), 7.30 (dd, ^*3*^*J*_*H12-H13*_ = 7.2 Hz, ^*3*^*J*_*H12-H11*_ = 4.2 Hz, 1H, **H**_**12**_).

^**13**^**C-NMR** (151 MHz, D_2_O, extracted from HSQC and HMBC, *δ*/ppm): 162.9 (**C**_**11**_), 154.4 (**C**_**4**_), 154.1 (**C**_**6**_), 149.9 (**C**_**2**_), 146.0 (**C**_**8**_), 138.4 (**C**_**13**_), 116.7 (**C**_**5**_), 112.0 (**C**_**12**_).

**HR-ESI-MS**: calcd. for [M+H]^+^ C_8_H_6_N_5_O m/z = 188.0567, found 188.0571.

#### Synthesis of N^6^-(3-Oxo-1-propenyl)-2’-deoxyadenosine (OPdA, CAS 178427-43-5)

OPdA was synthesized as previously described (*82*) with some modifications. 2 ′ - deoxyadenosine monohydrate (219 mg, 0.813 mmol, 1 eq., Sigma-Aldrich, Cat#D7400) was dissolved in 2 mL anhydrous dimethyl sulfoxide under argon atmosphere. Propargyl aldehyde (12 µl, 11.0 mg, 0.203 mmol, 0.25 eq., Toronto Research Chemicals, Cat#TRCP838440) was added to the stirred solution and additional propargyl aldehyde (1.25 eq.) was added over a 72-h period. The reaction mixture was filtered and purified by preparative HPLC on a Shimadzu LC system (LC-20AT prominence liquid chromatograph, with an SPD-20A prominence UV/VIS detector (λ = 254 and 280 nm). Preparative HPLC purification was performed using a Reprosil-Pur 120 ODS 3,5 µM, 150 × 20 mm column (Maisch GmbH, Cat#r15.93.s1520), where the mobile phases A and B were water and 90% acetonitrile in water, respectively. Separation was performed with a flow rate of 9 mL/min with a linear gradient of 1-30% B from 5 to 15 min, 30-100% B from 15 to 17 min, 100% B from 17 to 21 min, 100-0% B from 21 to 22 min and 1% B until 25 min. Analytical HPLC was performed with a LC-20AD prominence liquid chromatograph combined with a Shimadzu LCMS-2020 liquid chromatograph mass spectrometer. Biologically active HPLC peaks were collected for mass spectrometric and NMR analyses. The OPdA yield was 13.5 mg (44.0 μmol, 5.4%).

^**1**^**H-NMR** (500 MHz, D_2_O, *δ*/ppm): 9.21 (d, ^*3*^*J*_*H13-H12*_ = 8.7 Hz, 1H, **H**_**13**_), 8.49 (d, ^*3*^*J*_*H11-H12*_ = 13.5 Hz, 1H, **H**_**11**_), 8.43 (s, 1H, **H**_**8**_), 8.38 (s, 1H, **H**_**2**_), 6.45 (dd, ^*3*^*J*_*H1’-H2’a*_ = 6.8 Hz, ^*3*^*J*_*H1’-H2’b*_ = 6.8 Hz, 1H, **H**_**1’**_), 5.89 (dd, ^*3*^*J*_*H12-H11*_ = 13.5, ^*3*^*J*_*H12-H13* =_ 8.7 Hz, 1H, **H**_**12**_), 4.65 (ddd, ^*3*^J_*H3’-H2’a*_ = 6.1 Hz, ^*3*^J_*H3’-H2’b*_ = 3.5 Hz, ^*3*^J_*H3’-H4’*_ = 3.5 Hz, 1H, **H**_**3’**_), 4.19 (ddd, ^*3*^*J*_*H4’-H5’b*_ = 3.8 Hz, ^*3*^*J*_*H4’-H5’a*_ = 3.5 Hz, ^*3*^*J*_*H4’-H3’*_ = 3.5 Hz, 1H, **H**_**4’**_), 3.86 (dd, ^*2*^*J*_*H5’a-H5’b*_ = 12.5 Hz, ^*3*^*J*_*H5’a-H4’*_ = 3.4 Hz, 1H, **H**_**5’a**_), 3.80 (dd, ^*2*^*J*_*H5’b-H5’a*_ = 12.6 Hz, ^*3*^*J*_*H5’b-H4’*_ = 4.3 Hz, 1H, **H**_**5’b**_), 2.80 (ddd, ^*2*^*J*_*H2’a-H2’b*_ = 13.7 Hz, ^*3*^*J*_*H2’a-H1’*_ = 7.1 Hz, ^*3*^*J*_*H2’a-H3’*_ = 6.4 Hz, 1H, **H**_**2’a**_), 2.58 (ddd, ^*2*^*J*_*H2’b-H2’a*_ = 14.0 Hz, ^*3*^*J*_*H2’b-H1’*_ = 6.3 Hz, ^*3*^*J*_*H2’b-H3’ =*_ 3.5 Hz, 1H, **H**_**2’b**_).

^**13**^**C-NMR** (126 MHz, D_2_O, extracted from HSQC and HMBC, *δ*/ppm): 195.7 (**C**_**13**_), 151.9 (**C**_**2**_), 151.1 (**C**_**11**_), 150.6 (**C**_**4**_), 148.9 (**C**_**6**_), 142.6 (**C**_**8**_), 120.8 (**C**_**5**_), 111.1 (**C**_**12**_), 87.5 (**C**_**4’**_), 84.7 (**C**_**1’**_), 71.1 (**C**_**3’**_), 61.6 (**C**_**5’**_), 39.1 (**C**_**2’**_).

**HR-ESI-MS**: calcd. for [M+Na]^+^ C_13_H_15_N_5_NaO_4_ m/z = 328.1016, found 328.1020.

#### Synthesis of N^4^-(3-Oxo-1-propenyl)-2’-deoxycytidine (OPdC, CAS 129124-79-4)

OPdC was synthesized as previously described (*82*) with some modifications. 2 ′ - deoxycytidine (185 mg, 0.813 mmol, 1 eq., Sigma-Aldrich, Cat#D3897) was dissolved in 2 mL anhydrous dimethyl sulfoxide under argon atmosphere. Propargyl aldehyde (12.0 µL, 11.0 mg, 0.203 mmol, 0.25 eq.) was added to the stirred solution and additional propargyl aldehyde (1.25 eq.) was added over a 72-h period.

HPLC purification was performed as described for OPdA. The OPdC yield was 7.00 mg (25.0 μmol, 3.1%). Biologically active HPLC peaks were collected for mass spectrometric and NMR analyses.

^**1**^**H-NMR** (500 MHz, D_2_O, *δ*/ppm): 9.31 (d, ^*3*^*J*_*H10-H9*_ = 8.6 Hz, 1H, **H**_**10**_), 8.29 (d, ^*3*^*J*_*H8-H9*_ = 13.7 Hz, 1H, **H**_**8**_), 8.19 (d, ^*3*^*J*_*H6-H5*_ = 7.4 Hz, 1H, **H**_**6**_), 6.28 (d, ^*3*^*J*_*H5-H6*_ = 7.4 Hz, 1H, **H**_**5**_), 6.24 (dd, ^*3*^*J*_*H1’-H2’a*_ = 6.1 Hz, ^*3*^*J*_*H1’-H2’b*_ = 6.1 Hz, 1H, **H**_**1’**_), 5.91 (dd, ^*3*^*J*_*H9-H8*_ = 13.7 Hz, ^*3*^*J*_*H9-H10*_ = 8.6 Hz, 1H, **H**_**9**_), 4.43 (ddd, ^*3*^*J*_*H3’-H2’a*_ = 6.4 Hz, ^*3*^*J*_*H3’-H2’b*_ = 4.3 Hz, ^*3*^*J*_*H3’-H4’*_ = 4.3 Hz, 1H, **H**_**3’**_), 4.12 (ddd, ^*3*^*J*_*H4’-H5’b*_ = 4.8 Hz, ^*3*^*J*_*H4’-H5’a*_ = 4.1 Hz, ^*3*^*J*_*H4’-H3’*_ = 4.1 Hz, 1H, **H**_**4’**_), 3.87 (dd, ^*2*^*J*_*H5’a-H5’b*_ = 12.5 Hz, ^*3*^*J*_*H5’a-H4’*_ = 3.5 Hz, 1H, **H**_**5’a**_), 3.77 (dd, ^*2*^*J*_*H5’b-H5’a*_ = 12.5 Hz, ^*3*^*J*_*H5’b-H4’*_ = 5.3 Hz, 1H, **H**_**5’b**_), 2.55 (ddd, ^*2*^*J*_*H2’b-H2’a*_ = 14.1 Hz, ^*3*^*J*_*H2’b-H1’*_ = 6.3 Hz, ^*3*^*J*_*H2’b-H3’*_ = 4.3 Hz, 1H, **H**_**2’b**_), 2.32 (ddd, ^*2*^*J*_*H2’a-H2’b*_ = 14.2 Hz, ^*3*^*J*_*H2’a-H1’*_ = 6.5 Hz, ^*3*^*J*_*H2’a-H3’*_ = 6.5 Hz, 1H, **H**_**2’a**_).

^**13**^**C-NMR** (126 MHz, D_2_O, extracted from HSQC and HMBC, *δ*/ppm): 196.2 (**C**_**10**_), 161.5 (**C**_**4**_), 156.8 (**C**_**2**_), 149.6 (**C**_**8**_), 144.3 (**C**_**6**_), 112.1 (**C**_**9**_), 96.8 (**C**_**5**_), 87.1 (**C**_**1’**_), 87.1 (**C**_**4’**_), 70.3 (**C**_**3’**_), 61.1 (**C**_**5’**_), 39.8 (**C**_**2’**_).

**HR-ESI-MS**: calcd. for [M+Na]^+^ C_12_H_15_N_3_NaO_5_ m/z = 304.0904, found 304.0902.

### MS and NMR analysis

Unless otherwise stated, chemicals were used as received without further purification. NMR analysis of all of the antigens was performed at 298 K on a Bruker Avance III NMR spectrometer operating at 500 MHz proton frequency equipped with a BBFO probehead or on a Bruker Avance III HD NMR spectrometer operating at 600 MHz proton frequency equipped with a cryogenic QCI-F probe. Standard pulse sequences were used for cosy, tocsy, noesy, hsqc, hmqc and hmbc 2D-NMR experiments and the spectra were processed using the topspin 4.0 software package. All compounds were fully characterized by means of 2D-NMR and HR-ESI-MS. For all compounds ^1^H- and ^1^H-^13^C-HSQC or ^1^H-^13^C-HMQC spectra, as well as experimental and calculated HR-ESI-MS spectra are shown in Figure S3 and S4.

HR-ESI-MS spectra were measured on a Bruker MaXis 4G high resolution ESI Mass Spectrometer in direct injection mode using methanol containing 0.1% v/v formic acid.

### Evaluation of compound biological activity

#### Cell surface MR1 upregulation

THP-1 cells (10^5^ cells/well) were tested for MR1 surface expression after incubation with or without synthetic compounds: OPdA (100 μM), M_1_G (13μM) and OPdC (100 μM) for 6 h at 37°C. Ac-6-FP (100 µM) (Schircks Laboratories, Cat#11.418) was used as positive control for MR1 surface upregulation. The cells were stained with an anti-human-MR1-APC mAb (clone 26.5, Biolegend) or with APC-labeled mouse IgG2a, k isotype control (clone MOPC-173, Biolegend) antibodies for 20 min at 4°C, then washed and analyzed by flow cytometry. For each condition, net MFI was calculated subtracting isotype MFI from anti-MR1 MFI and fold change of cells treated with synthetic molecules over cells treated with vehicle was calculated.

#### T-cell activation assays with living or fixed APCs

MR1T cells (5×10^4^/well unless otherwise indicated) were co-cultured with the indicated APCs (10^5^ cells per well unless otherwise indicated) for 18 h in 120 µL volume in triplicate. In some experiments, anti-MR1 mAbs (clone 26.5, Ultra-LEAF, Biolegend, Cat#361110) or mouse IgG2a isotype control mAbs (clone MOPC-173, Ultra-LEAF, Biolegend, Cat#400263) (both at 30 µg/mL) were added and incubated for 30 min at 37°C prior to the addition of T cells.

When nucleobases, nucleosides or nucleotides (all 250 µM, all from Sigma-Aldrich) and synthetic compounds OPdA, M_1_G or OPdC were used to stimulate T cells, the THP-1 cells (10^5^/well) were cultured 2 h with the indicated molecules or medium only, prior to T-cell addition. When a single concentration of synthetic antigens was used, 100 µM OPdA, 100 µM OPdC and 13 µM M_1_G were used for all clones.

In experiments with Mycophenolic acid (10 µM, Sigma-Aldrich, Cat#M3536), EHNA (25 µM, Sigma-Aldrich, Cat#E114), S-p-bromobenzylglutathione cyclopentyl diester (BBG, 20 µM, Sigma-Aldrich, Cat#SML1306), THP-1 cells (1×10^6^/mL) were treated with the indicated concentrations of drugs in complete medium at 37°C for 18 h before being washed twice with PBS, counted and used for T-cell activation. In experiments where THP-1 cells (1×10^6^/mL) were treated with Doxorubicin (75 nM, Selleck Chemical, Cat#S1208) and Paclitaxel (5 µM, Sigma-Aldrich, Cat#T7191), the cells were incubated at 37°C for 18 h before being washed twice with PBS, counted, and incubated for 2 h with vehicle or 150 µM dAdo (Sigma-Aldrich Cat#D7400) and 150 µM Guanosine (Sigma-Aldrich, Cat#G6752) prior to T-cell addition. In experiments where fixed A375-MR1 cells were used to activate MR1T cells, APCs (4×10^5^ /mL) were treated with Apocynin (APO, 100 µM, Cat#W508454), L-Glutathione reduced (GSH, 4 mM, Cat#G6013), N-acetyl cysteine (NAC, 4 mM, Cat#A9165), L-Buthionine-sulfoximine (BSO, 400 µM, Cat#B2515) Mercaptosuccinic acid (MSA, 3.3 µM, Cat#M6182), ML-210 (6 µM, Cat#SML0521) or 1S,3R-RSL3 (RSL3, 1 µM, Cat#SML2234), Hydralazine hydrochloride (100 µM, Cat#H1753) or Aminoguanidine hemisulfate salt (5 mM, Cat#A7009) for 18 h at 37°C before being washed twice with PBS, fixed with Glutaraldehyde (Cat#G5882), counted and used for MR1T cell stimulation (all drugs are from Sigma-Aldrich).

In some experiments, MAIT cells were stimulated by APCs pulsed 3 h with 5-OP-RU as previously described (*83*) or with 30 µM 6,7-dimethyl-8-Ribityllumazine (Cayman Chemical Cat#23370).

To confirm that drugs reducing MR1T stimulation do not affect MR1 presentation ability, A375-MR1 cells treated with different molecules were collected before fixation and used to stimulate the MAIT clone MRC25 after pulsing for 2 h at 37°C with the indicated concentrations of freshly-prepared 5-OP-RU (*83*) or 6,7-dimethyl-8-Ribityllumazine.

#### Activation assay with plate-bound soluble MR1

Recombinant human β2m-MR1-K43A-Fc was produced in CHO-K1 cells as previously described (*10*) and 4 µg/mL were coated onto 96 wells plates (Nunc, Cat#439454) for 18 h at 4°C. Plate-bound MR1 was then washed twice with wash buffer (150 mM NaCl, 20 mM Tris and 2% Glycerol, pH 5.6) to remove bound antigens. Then, the synthetic antigens (OPdA, M_1_G and OPdC) were added at the indicated concentrations and incubated for 6 h at room temperature. Unbound antigens were washed twice with PBS before the addition of excess PBS. Indicated MR1T cell clones (10^5^/100 µl/well) were added and supernatants were collected after 18 h. Released cytokines were detected by ELISA.

### Cytokine analysis

The following human cytokines were assessed by ELISA as previously described (*10*): human IFN-γ (capture MD-1 mAb; revealing biotinylated 4S.B3 mAb, Biolegend Cat#507502 and 502504, respectively), human IL-13 (capture JES10-5A2 mAb; revealing biotinylated SB126d mAb, SouthernBiotech Cat#10125-01 and 15930-08, respectively).

### Immunofluorescence staining

Cell surface labeling was performed using standard protocols and all mAbs were titrated on appropriate cells before use. The samples were acquired on an LSR Fortessa flow cytometer with the FACS Diva software (Becton Dickinson) or on CytoFLEX with CytExpert software (Beckman Coulter). Cell sorting experiments were performed using either Influx or FACSaria (Becton Dickinson). Dead cells and doublets were excluded on the basis of forward scatter area and width, side scatter, and DAPI (Sigma-Aldrich, Cat#D9542) or Live/Dead DRAQ7™ (Thermo Fisher Scientific Cat#424001) staining. All data were analyzed using FlowJo (v10.5.3 LLC).

### ROS production measurement

CM-H_2_DCFDA (Thermo Fisher Scientific, Cat#C6827) was used to assess ROS production in cell upon cell treatment with Doxorubicin and Paclitaxel. THP-1 cells (10^7^/mL) were labelled with 10 µM CM-H_2_DCFDA for 30 min at 37°C in the dark, then washed with PBS and resuspended in complete medium. 10^5^ cells were seeded per well and treated with 75 nM Doxorubicin, 5 µM Paclitaxel or vehicle for 18 h at 37°C. Phorbol 12-myristate 13-acetate (PMA, 50 ng/mL, Sigma-Aldrich, Cat#P8139) was used as positive control.

## QUANTIFICATION AND STATISTICAL ANALYSIS

### sgRNA-Seq data processing and analysis

Raw sequencing data was demultiplexed using bcl2fastq (v2.17.1.14, Illumina) and read quality checked using FastQC (v0.11.4, Babraham Bioinformatics v0.11.4). Reads were then trimmed to remove the homologous regions flanking the sgRNA sequences using Trimmomatic (v0.36, http://www.usadellab.org/cms/?page=trimmomatic) (*84*) using options HEADCROP:42 CROP:20. These trimmed reads were again passed through FastQC to check that average phred33 quality in the sgRNA sequences was >30. These reads were aligned to a GeCKO v2 sgRNA reference index using Bowtie2 (v2.2.9, http://bowtie-bio.sourceforge.net/bowtie2/index.shtml) (*85*) with options --very-sensitive-local. Read counts were then extracted from the resulting SAM files using custom perl script map_count.pl (Cox, M. available on request) and imported into R (*86*) for analysis using edgeR (v3.24.3, www.bioconductor.org) (*87*).

Inequality of sgRNA activity in the GeCKO library (*88*) hinders hit-selection using rank-based methodology, thus differential enrichment analysis was performed with the edgeR package (*87*) using the GLM Robust method to estimate dispersions, after removing guides targeting known essential genes (*16*). The level of random enrichment and depletion of guides was estimated using the log_2_ fold-change in the top and bottom 1% of negative control guides in the GeCKO library. Thus, guides with an FDR < 0.05 and log_2_ fold-change greater or lower than the top and bottom 1% of negative control guides, respectively, were said to be significantly enriched or depleted by our screen. GO-term and KEGG-pathway enrichment analysis was performed using a binomial test on the significant unique gene-targets identified in the differential enrichment analysis (GO.db, v3.7.0)(*89*). Genes were annotated using biomaRt (v2.42.0, www.bioconductor.org) (*90*).

### Metabolic modeling

To perform structural sensitivity analysis (*21, 91*), the Recon3D model was pre-processed by making all reactions reversible and removing biomass, maintenance, sink, and demand reactions. For each gene in the model, associated reactions were aligned by reversing reaction directions such that the overlap of metabolites on each side of the reaction equations was maximal. Assuming that perturbation of a gene implied identical flux perturbations for all associated reactions, flux balance analysis (FBA) (*92*) was used to check if this identical flux perturbation was feasible at steady state. If it was infeasible, the closest feasible perturbation was found by minimizing the Euclidean distance to the identical flux perturbation. Finally, perturbed fluxes were fixed and the minimal network response was computed by minimizing the Euclidean norm of all fluxes not directly linked to the perturbed gene. If the minimal response was a thermodynamically infeasible loop (*93*), constraints and variables were added to specifically disable this loop (*94*) and a minimal response was re-computed until the minimal loopless response was found. Genome-wide knock-out sensitivities were analyzed in R to identify sets of >2 knock-outs with Pearson correlation scores > 0.6.

### Data analysis and statistical analysis

Data analysis, statistical tests and visualization were conducted in R and GraphPad Prism. In Prism (v8.1.0, GraphPad Software, Inc.), analysis was performed using multiple t-test, One- or Two-way Anova as indicated for each assay in figure legends.

A p value < 0.05 was considered statistically significant. *p < 0.05, **p ≤ 0.01, ***p ≤ 0.005.

## Data availability

sgRNA sequencing data have been deposited to the NCBI GEO repository accession GSE160366.

