## Supplemental Information for "Nucleobase adduct-containing metabolites are MR1 ligands that stimulate self-reactive MR1T cells"

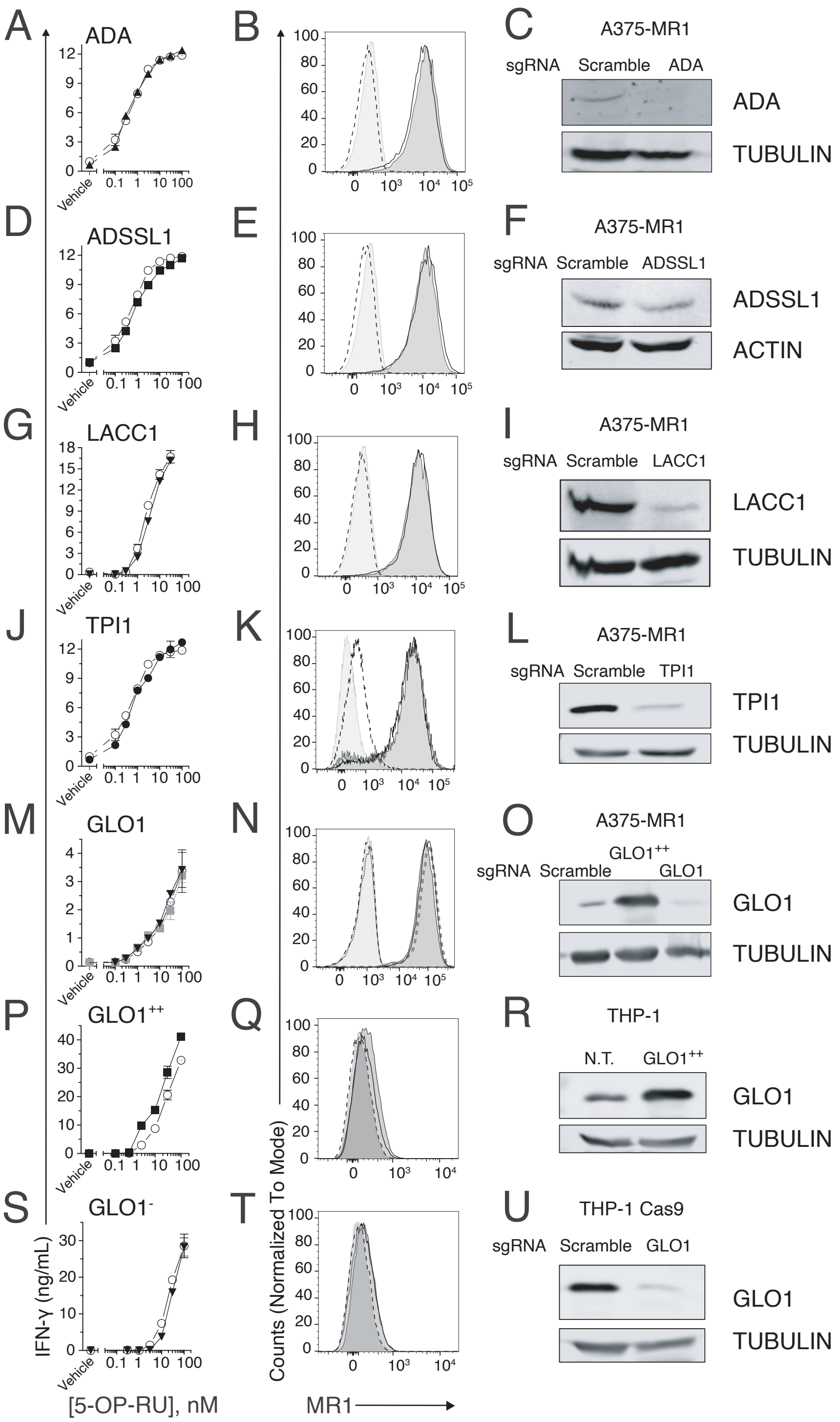


**Figure S1 Genetically-engineered cell lines characterization, Related to Figures 2 and 3.**

(A, D, G, J, M, P, S) Activation assay of MAIT clone MRC25 in response to A375-MR1 (A, D, G, J, M) and THP-1 (P and S) cells pulsed with 5-OP-RU. Cells are either wild type (○), knock-out (A ▲, D ■, G ▼, J ●, M ▼, S ▼) or overexpressing (M ■, P ■) the indicated genes. IFN-γ is expressed as mean ± SD of triplicate independent cultures.

(B, E, H, K, N, Q, T) Surface MR1 expression of the genetic engineered cell lines. MR1 staining of wild type cells (dark grey shadow), ko lines (black line) and GLO1-overexpressing A375-MR1 (N, grey thick dashed line) with anti-MR1 mAbs 26.5. Isotype-matched control staining is depicted in wild type cells (light grey shadow with grey dot line), in ko cells (black dashed line) or GLO1-overexpressing A375-MR1 (N, black dotted line).

(C, F, I, L, O, R, U) Western blot analysis of target protein expression in indicated cell lines. Tubulin or Actin were used as loading control.

The experiments were repeated at least twice and one representative experiment is shown.

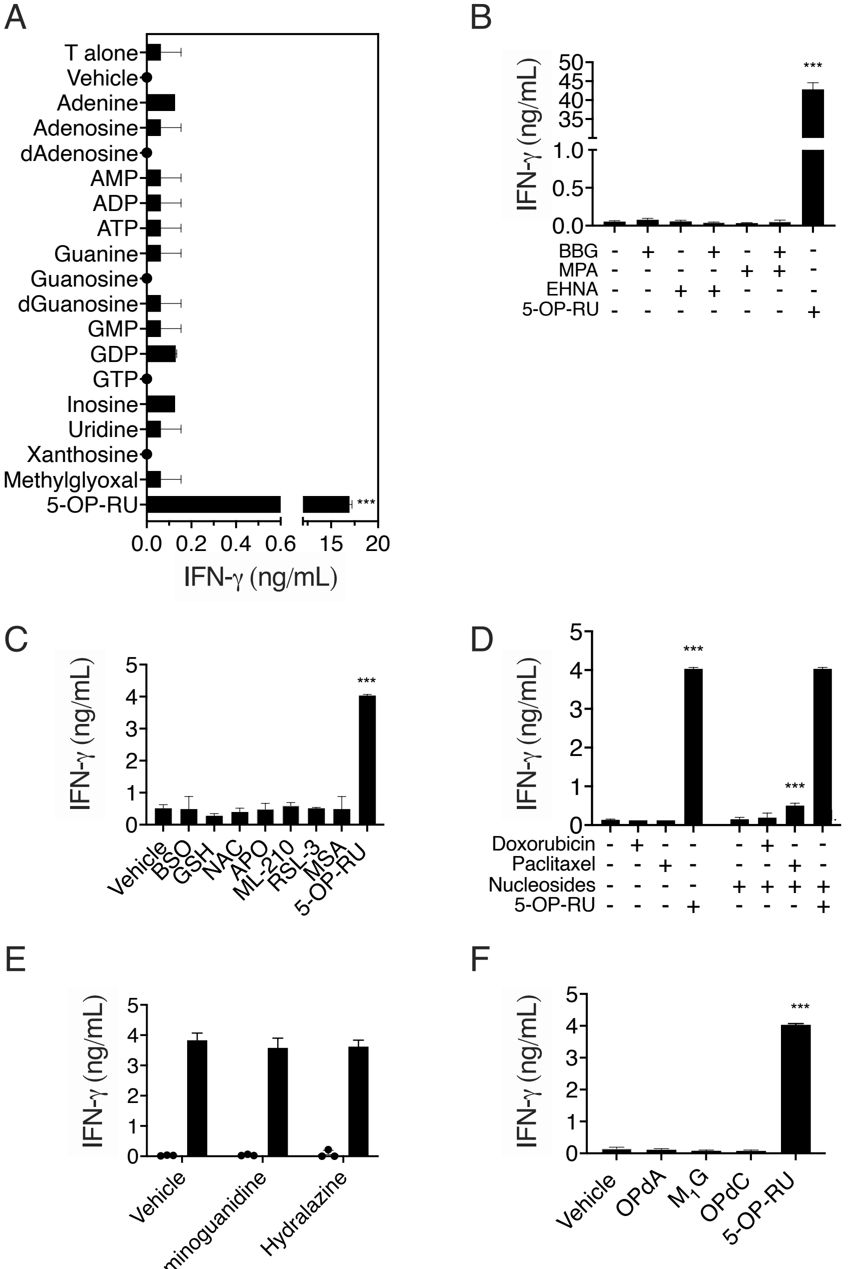


**Figure S2. Stimulation of MAIT clone MRC25 with nucleobases, inhibitory drugs and synthetic antigens, Related to Figures 3, 4, 5**

(A) MAIT clone MRC25 was stimulated with THP-1 cells in the presence of different nucleobases (250 µM), Methylglyoxal (250 µM) or 5-OP-RU (30 nM).

(B and D) MRC25 cells were stimulated with THP-1 cells treated with indicated drugs.

(C) MRC25 cells were stimulated with A375-MR1 cells treated with GSH, NAC, APO, BSO or GPX inhibitors and fixed or with THP-1 cells pulsed with 5-OP-RU (10 nM).

(E) MRC25 cells were stimulated with A375-MR1 cells treated with carbonyl scavengers at indicated concentrations and fixed before T cell addition (empty bars). As controls, the same experiment was performed with the same carbonyl scavengers in the presence of 6,7-dimethyl-8-Ribityllumazine (20 µM, black bars).

(F) MRC25 cells were stimulated with THP-1 cells in the presence of OPdA, OPdC (both 100 µM), $M_1G$ (13 µM) or 5-OP-RU (10 nM).

n.d.=not determined

***p< 0.001 compared to vehicle-treated cells using One-way ANOVA (A, B, C, F) or Two-way ANOVA (D and E) with Dunnett's multiple comparison.

IFN-γ is expressed as mean ± SD of triplicate independent cultures. The experiments were repeated at least twice and one representative experiment is shown.

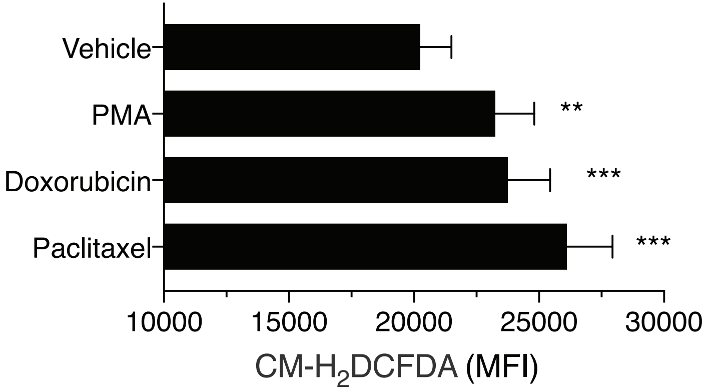


**Figure S3 ROS accumulation in APCs, Related to Figures 4**
Quantification of ROS produced in THP-1 cells treated with Doxorubicin, Paclitaxel or Phorbol 12-myristate 13-acetate (PMA). Results are expressed Median Fluorescence Intensity (MFI) of live cells ± SD. Experiment was performed in technical triplicates, was repeated twice and one representative experiment is shown.
**$p \leq 0.01$ and ***$p \leq 0.001$ using one-way Anova with Dunnett's multiple comparison

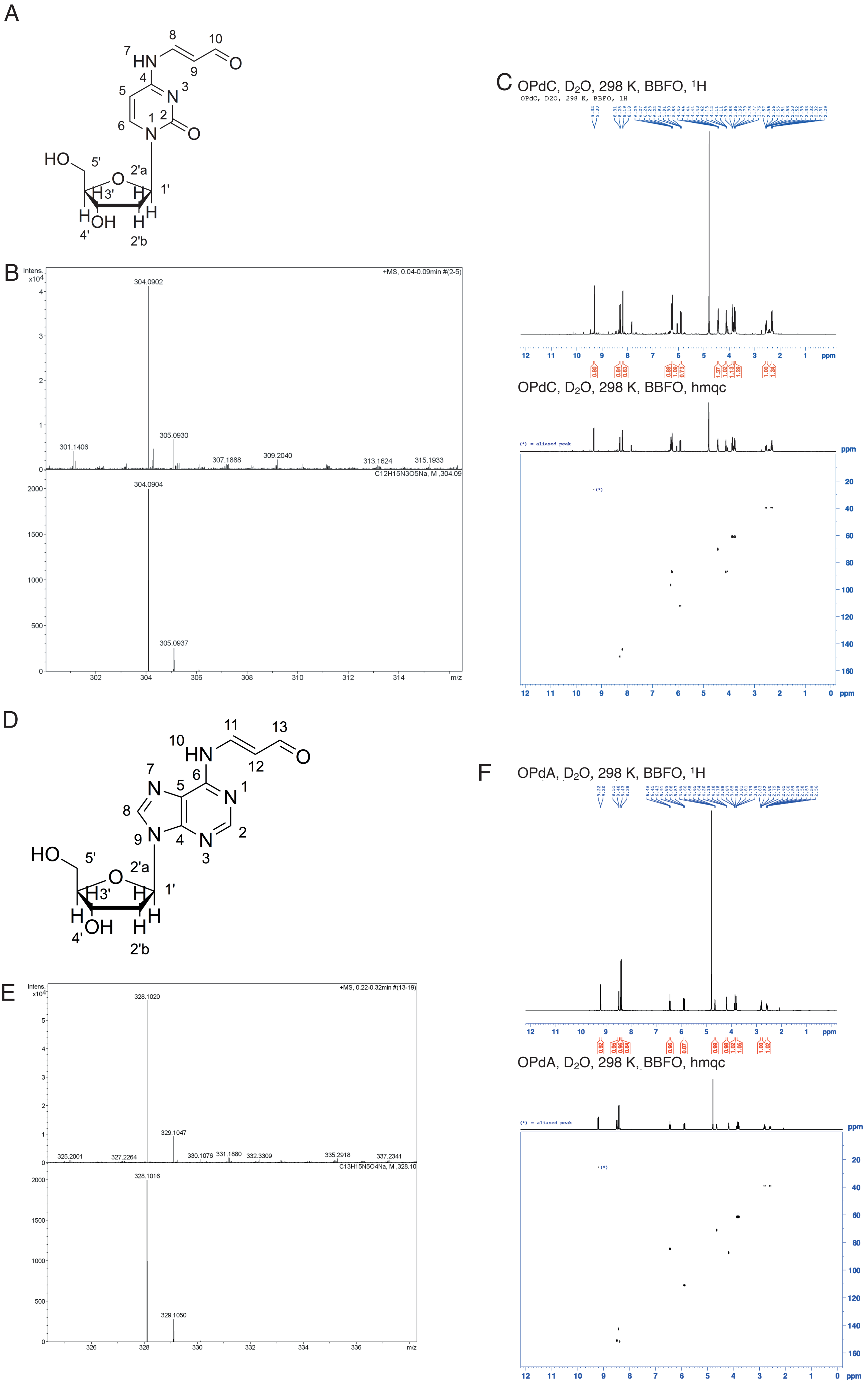


**Figure S4. Mass-Spectrometric and NMR analyses of synthetic OPdC and OPdA, Related to Figure 5.**
(A) $N^4$-(3-Oxo-1-propenyl)-2'-deoxycytidine (OPdC) structure, (B) HRMS, top: experiment, bottom: calculated, and (C) NMR analysis.
(D) $N^6$-(3-Oxo-1-propenyl)-2'-deoxyadenosine (OPdA) structure, (E) HRMS, top: experiment, bottom: calculated, and (C) NMR analysis.

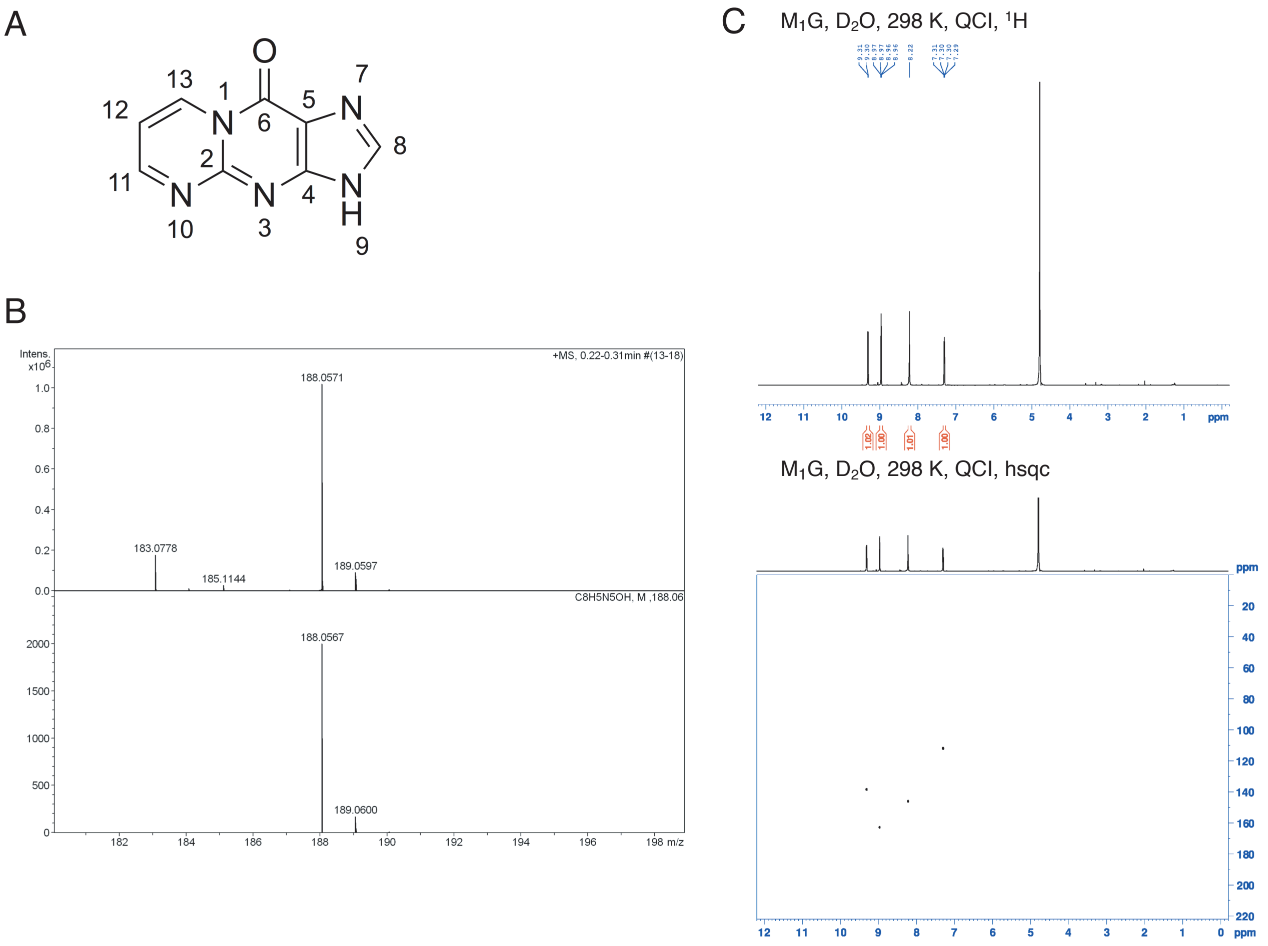


**Figure S5. Mass-Spectrometric and NMR analyses of synthetic $M_1G$. Related to Figure 5.**
(A) Pyrimido[1,2- $\alpha$ ] purin-10(3*H*)-one ($M_1G$) structure, (B) HRMS, top: experiment, bottom: calculated, and (C) NMR analysis.

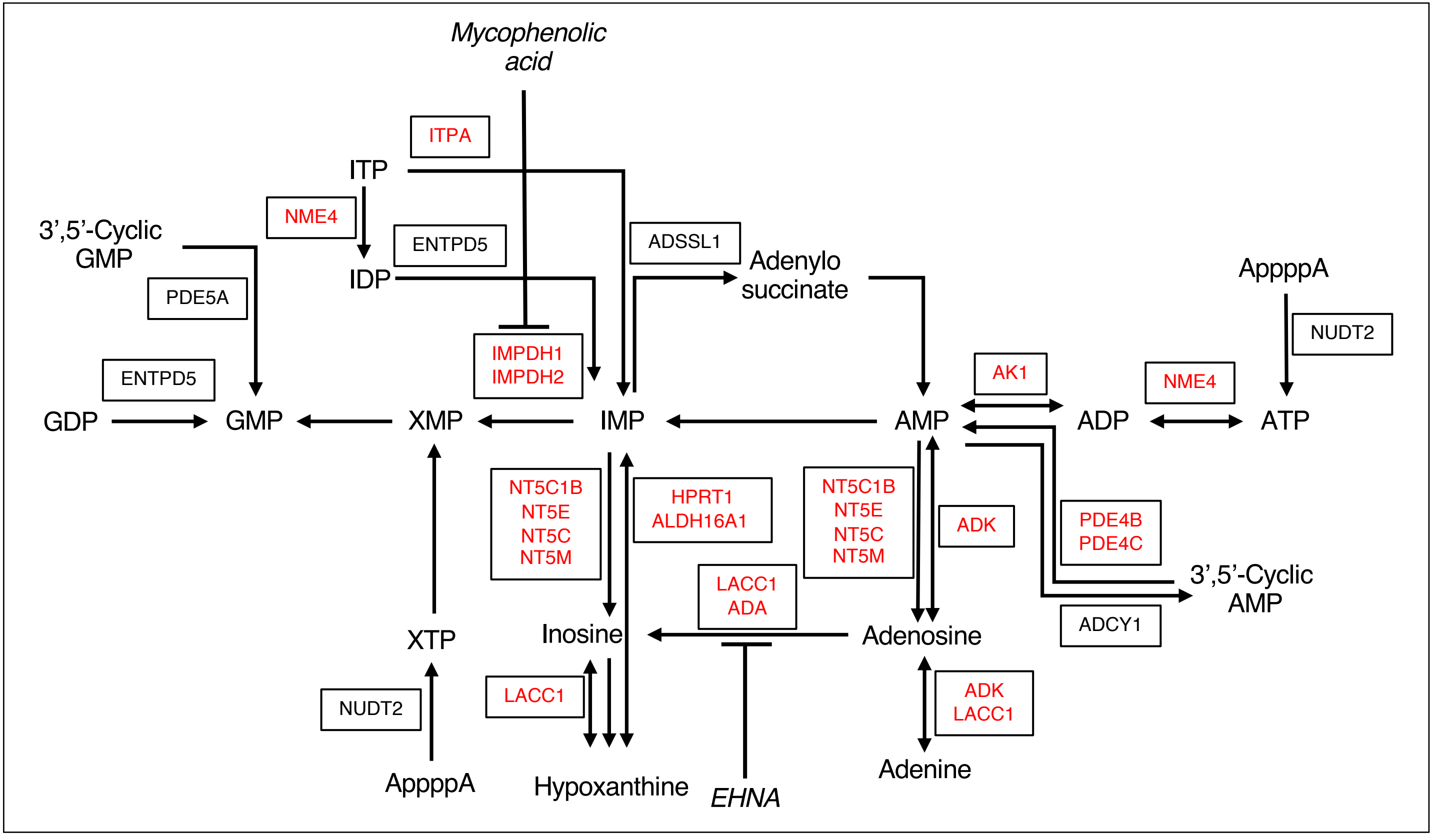


**Scheme S1 sgRNA hits identified in purine metabolism pathway, Related to Figures 1-3 and Data S1 and S2.** Simplified representation of the purine metabolism pathway with significantly enriched (black) or depleted (red) sgRNA targets found in the genome-wide CRISPR/Cas9 screening. Drugs used to inhibit specific enzymes are *italicized*.

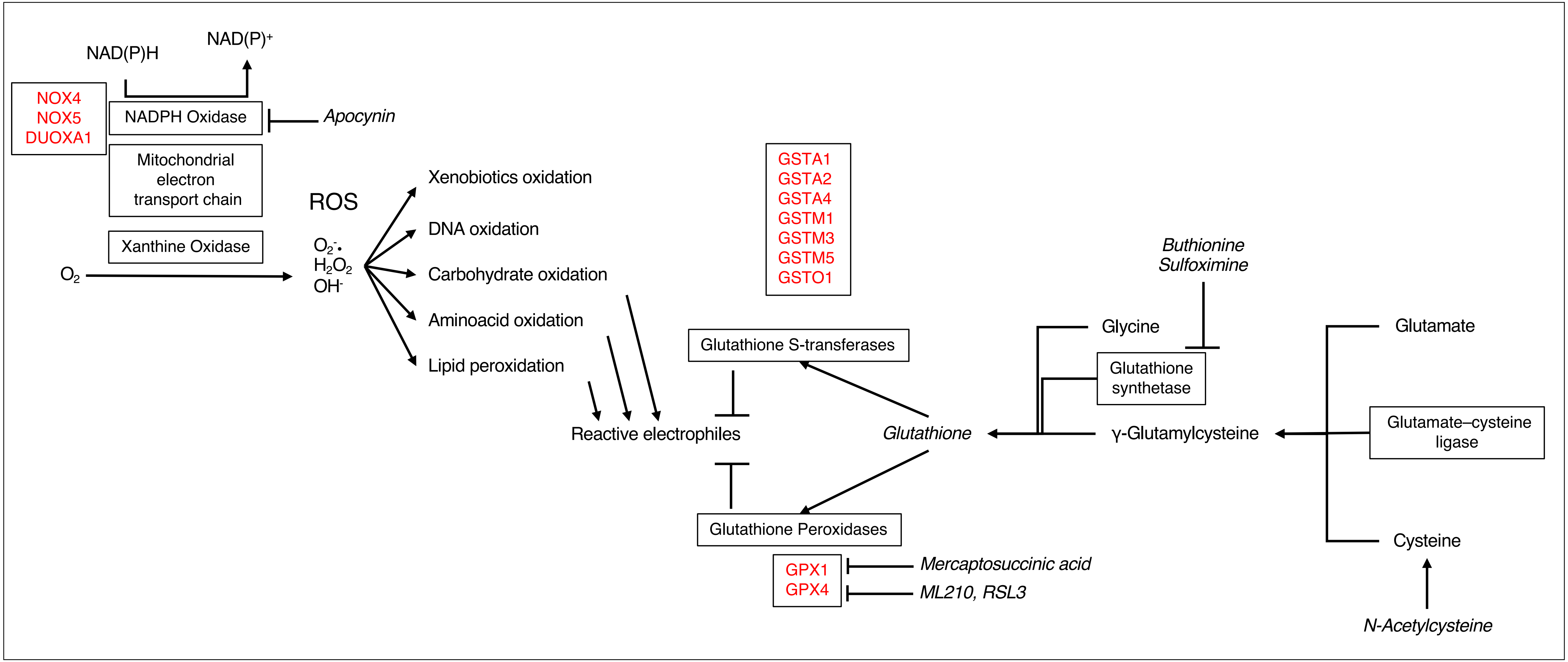


**Scheme S2** ROS generation and scavenging, Related to Figure 4 and Data S1 and S2.
Simplified representation of ROS production and scavenging pathways. Significantly depleted sgRNA targets found in the genome-wide CRISPR/Cas9 screening are annotated in red. Drugs and compounds used are *italicized*.

| Kegg Pathway | KEGG ID | Freq | Genome-wide Ratio | Sample Ratio | p-value | FDR | q-value |
|---|---|---|---|---|---|---|---|
| **Enriched Pathways** | | | | | | | |
| Purine metabolism | hsa00230 | 5 | 0.060341821 | 0.263157895 | 0.004552013 | 0.022760067 | ND |
| Mannose type O-glycan biosynthesis | hsa00515 | 2 | 0.007673526 | 0.105263158 | 0.009232434 | 0.023081085 | ND |
| **Depleted Pathways** | | | | | | | |
| Glycolysis / Gluconeogenesis | hsa00010 | 19 | 0.023369376 | 0.049608355 | 0.002005235 | 0.033874609 | 0.01353054 |
| Fatty acid degradation | hsa00071 | 12 | 0.014998256 | 0.031331593 | 0.014214189 | 0.067458054 | 0.02478919 |
| Oxidative phosphorylation | hsa00190 | 25 | 0.041855598 | 0.065274151 | 0.020345478 | 0.067458054 | 0.02478919 |
| Purine metabolism | hsa00230 | 36 | 0.060341821 | 0.093994778 | 0.006131678 | 0.047124933 | 0.01592725 |
| Glutathione metabolism | hsa00480 | 14 | 0.019532612 | 0.036553525 | 0.020048346 | 0.067458054 | 0.02478919 |
| Arachidonic acid metabolism | hsa00590 | 18 | 0.021276596 | 0.046997389 | 0.001710169 | 0.033874609 | 0.01353054 |
| Metabolism of xenobiotics by cytochrome P450 | hsa00980 | 17 | 0.025113359 | 0.044386423 | 0.018309849 | 0.067458054 | 0.02478919 |
| Drug metabolism - cytochrome P450 | hsa00982 | 18 | 0.023718172 | 0.046997389 | 0.005251754 | 0.047124933 | 0.01592725 |

**Table S1. KEGG pathways related to statistically significant enriched or depleted sgRNAs.**

**Related to Figure 1, Data S1 and S2.** Frequency refers to the number of unique significantly enriched/depleted gene-targets annotated in each pathway listed. The Genome-wide and Sample ratios are the proportion of genes within the human genome and unique significantly enriched/depleted gene-targets annotated within each pathway, respectively. Binomial enrichment testing provided p-values indicating most common pathways. Multiple testing correction was performed by computing the Benjamini-Hochberg false discovery rate (FDR) and q-value.

| Target gene | gRNA sequence | Vector | Source |
|---|---|---|---|
| *ADA* | CAGGCTTGATGGATCCGTCT | pLV-mCherry-U6 gRNA | VectorBuilder |
| *ADA* | TCACCGTACTGTCCACGCCG | lentiGuide-Puro | This paper |
| *ADA* | GCGGTACAGTCCGCACCTGC | lentiGuide-Puro | This paper |
| *ADSSL1* | TTCCAGGGGGGCAACAACGC | lentiGuide-Puro | This paper |
| *ADSSL1* | GCTGATGATGTCGGCGTCCG | lentiGuide-Puro | This paper |
| *ADSSL1* | ACATACCGAAGTCAATGTCG | lentiGuide-Puro | This paper |
| *LACC1* | CGTAGGTTGGCGAATGCTGC | lentiGuide-Puro | This paper |
| *LACC1* | TACCTTGGGATCTCTCCGTT | lentiGuide-Puro | This paper |
| *LACC1* | TCAAGAAAATCTGCGTAGGT | lentiGuide-Puro | This paper |
| *GLO1* | GAACCGCAGCCCCCGTCCGG | pRP[gRNA]-EGFP:P2A:Puro-U6 | VectorBuilder |
| *GLO1* | GTCCGGCGGCCTCACGGACG | pRP[gRNA]-EGFP:P2A:Puro-U6 | VectorBuilder |
| *TPI1* | CGGCGAGGGCTTACCGGTGT | lentiGuide-Puro | This paper |
| *TPI1* | ACCGGTGTCGGCCGGCACCT | lentiGuide-Puro | This paper |
| *TPI1* | CGAAGTCGATATAGGCAGTA | lentiGuide-Puro | This paper |
| Scrambled sequence | GTGTAGTTCGACCATTCGTG | lentiGuide-Puro | This paper |
| Scrambled sequence | GTTCAGGATCACGTTACCGC | lentiGuide-Puro | This paper |
| Scrambled sequence | AAATGTGAGATCAGAGTAAT | lentiGuide-Puro | This paper |

**Table S2. sgRNA target sequences used for *knock-out* generation, Related to Figures 2 and 3.**
lentiGuide-Puro, Addgene, Cat#352963.
